# Structural basis for divergent and convergent evolution of catalytic machineries in plant aromatic amino acid decarboxylase proteins

**DOI:** 10.1101/404970

**Authors:** Michael P. Torrens-Spence, Ying-Chih Chiang, Tyler Smith, Maria A. Vicent, Yi Wang, Jing-Ke Weng

## Abstract

Radiation of the plant pyridoxal 5’-phosphate (PLP)-dependent aromatic L-amino acid decarboxylase (AAAD) family has yielded an array of paralogous enzymes exhibiting divergent substrate preferences and catalytic mechanisms. Plant AAADs catalyze either the decarboxylation or decarboxylation-dependent oxidative deamination of aromatic L-amino acids to produce aromatic monoamines or aromatic acetaldehydes, respectively. These compounds serve as key precursors for the biosynthesis of several important classes of plant natural products, including indole alkaloids, benzylisoquinoline alkaloids, hydroxycinnamic acid amides, phenylacetaldehyde-derived floral volatiles, and tyrosol derivatives. Here, we present the crystal structures of four functionally distinct plant AAAD paralogs. Through structural and functional analyses, we identify variable structural features of the substrate-binding pocket that underlie the divergent evolution of substrate selectivity toward indole, phenyl, or hydroxyphenyl amino acids in plant AAADs. Moreover, we describe two mechanistic classes of independently arising mutations in AAAD paralogs leading to the convergent evolution of the derived aldehyde synthase activity. Applying knowledge learned from this study, we successfully engineered a shortened benzylisoquinoline alkaloid pathway to produce (S)-norcoclaurine in yeast. This work highlights the pliability of the AAAD fold that allows change of substrate selectivity and access to alternative catalytic mechanisms with only a few mutations.

**Significance:** Plants biosynthesize their own proteinogenic aromatic L-amino acids, namely L-phenylalanine, L-tyrosine and L-tryptophan, not only for building proteins but also for the production of a plethora of aromatic-amino-acid-derived natural products. Pyridoxal 5’-phosphate (PLP)-dependent aromatic L-amino acid decarboxylase (AAAD) family enzymes play important roles in channeling various aromatic L-amino acids into diverse downstream specialized metabolic pathways. Through comparative structural analysis of four functionally divergent plant AAAD proteins together with biochemical characterization and molecular dynamics simulations, we reveal the structural and mechanistic basis for the rich divergent and convergent evolutionary development within the plant AAAD family. Knowledge learned from this study aids our ability to engineer high-value aromatic-L-amino-acid-derived natural product biosynthesis in heterologous chassis organisms.

## Introduction

Plants are sessile organisms that produce a dazzling array of specialized metabolites as adaptive strategies to mitigate multitudes of abiotic and biotic stressors. Underlying plants’ remarkable chemodiversity is the rapid evolution of the requisite specialized metabolic enzymes and pathways (1). Genome-wide comparative analysis across major green plant lineages has revealed pervasive and progressive expansions of discrete enzyme families mostly involved in specialized metabolism, wherein new enzymes emerge predominantly through gene duplication followed by subfunctionalization or neofunctionalization (2). Newly evolved enzyme functions usually entail altered substrate specificity and/or product diversity without changes in the ancestral catalytic mechanism. In rare cases, adaptive mutations occur in a progenitor protein fold—either catalytic or non-catalytic—that give rise to new enzymatic mechanisms and novel biochemistry (3, 4). Resolving the structural and mechanistic bases for these cases is an important step towards understanding the origin and evolution of functionally disparate enzyme families.

Aromatic amino acid decarboxylases (AAADs) are an ancient group of pyridoxal 5’-phosphate (PLP)-dependent enzymes with primary functions associated with amino acid metabolism. Mammals possess a single AAAD, 3,4-dihydroxyphenylalanine decarboxylase (DDC), responsible for the synthesis of monoamine neurotransmitters, such as dopamine and serotonin from their respective aromatic L-amino acid precursors (5). In contrast, AAAD family enzymes have radiated in plants and insects to yield a number of paralogous enzymes with variation in both substrate preference and catalytic mechanism (Fig. S1) (6). In plants, L-tryptophan decarboxylases (TDCs) and L-tyrosine decarboxylases (TyDCs) are canonical AAADs that supply the aromatic monoamine precursors tryptamine and tyramine for the biosynthesis of monoterpene indole alkaloids (MIAs) and benzylisoquinoline alkaloids (BIAs), respectively (Fig. S2) (7, 8). In contrast, phenylacetaldehyde synthases (PAASs) and 4-hydroxyphenylacetaldehyde synthases (4HPAASs) are mechanistically divergent aromatic aldehyde synthases (AASs) that catalyze the decarboxylation-dependent oxidative deamination of L-phenylalanine and L-tyrosine, respectively, to produce the corresponding aromatic acetaldehydes necessary for the biosynthesis of floral volatiles and tyrosol-derived natural products (Fig. S2) (9, 10). Plant AAAD or AAS enzymes thus play a gate-keeping role in channeling primary metabolites into a variety of specialized metabolic pathways.

In this work, we seek to understand the evolutionary trajectories that led to the functional divergence and convergence within the plant AAAD family proteins. Through comparative analysis of four representative plant AAAD crystal structures followed by mutant characterization and molecular dynamics (MD) simulation, we identified key structural features that dictate substrate selectivity and the alternative AAAD-versus-AAS catalytic mechanisms. Using these findings, we discovered a group of myrtle plant AASs with a catalytic residue substitution similar to those described in insect AASs but distinct from the substitutions previously described in plant AASs (10). These findings suggest that nature has explored multiple mechanistically distinct trajectories to reconfigure an ancestral AAAD catalytic machinery to catalyze AAS chemistry. Furthermore, we show the feasibility of engineering a shortened BIA biosynthetic pathway in yeast *Saccharomyces cerevisiae* by harnessing the 4HPAAS activity, highlighting the role of a neofunctionalized enzyme in rewiring ancestral metabolic network during plant specialized metabolic evolution.

## Results

### The overall structures of four divergent plant AAAD proteins

To understand the structural basis for the functional divergence of plant AAAD proteins, we determined the X-ray crystal structures of *C. roseus* TDC (*Cr*TDC) in complex with L-tryptophan, *P. somniferum* TyDC (*Ps*TyDC) in complex with L-tyrosine, *A. thaliana* PAAS (*At*PAAS) in complex with L-phenylalanine, and unbound *R. rosea* 4HPAAS (*Rr*4HPAAS) (Fig. 2*A*, Fig. S3 and Table S1). All four enzymes pack in the crystal lattice as homodimers, and share the typical type II PLP-dependent decarboxylase fold (Table S2). Each monomer contains three distinct segments (Fig. S5), which were previously described as domains in the homologous mammalian DDC structure (5). These segments are unlikely to be stable as autonomously folding units, and rather function as stretches of topologically associated constituents necessary for the overall architecture of the α2 dimer. As represented by the L-tryptophan-bound *Cr*TDC structure, the N-terminal segment (*Cr*TDC^1-119^) comprises three antiparallel helices that interlock with the reciprocal helices of the other monomer to form the primary hydrophobic dimer interface (Fig. 2*B* and Fig. S6). The middle (*Cr*TDC^120-386^) segment harbors a conserved Lys^319^ with its ζ-amino group covalently linked to the coenzyme PLP (termed as LLP hereafter), and, together with the C-terminal (*Cr*TDC^387-497^) segment, forms two symmetric active sites at the dimer interface (Fig. 2*C*).

**Fig. 1.**
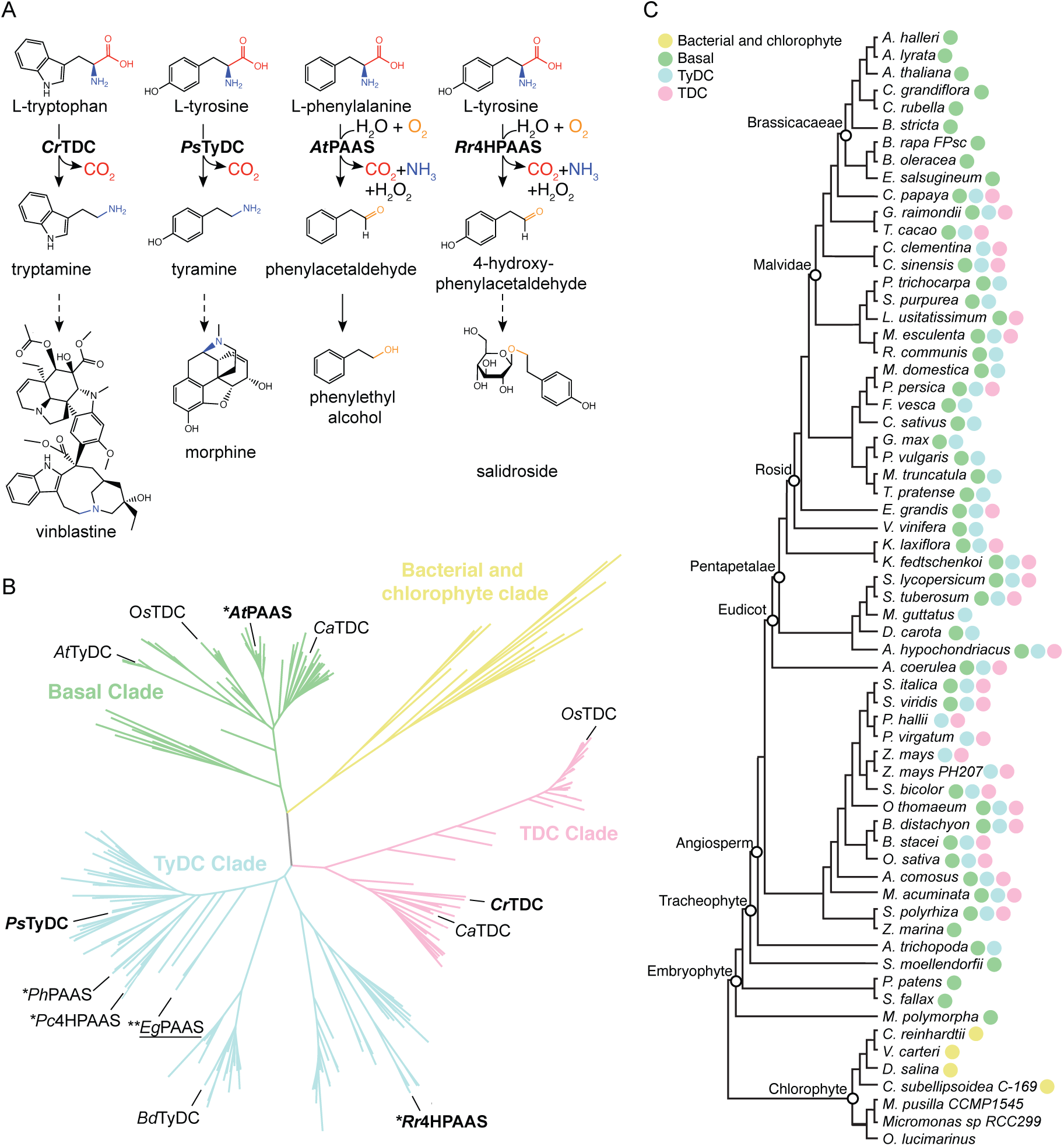
Function, phylogeny, and taxonomic distribution of plant AAADs. (*A*) Biochemical functions of four representative plant AAADs in the context of their native specialized metabolic pathways. The dotted arrows indicate multiple catalytic steps. (*B*) A simplified maximum-likelihood phylogenetic tree of bacteria, chlorophyte, and plant AAADs. A fully annotated tree is shown in Fig. S3. The bacterial/chlorophyte, basal, TyDC, and TDC clades are colored in yellow, green, blue, and pink, respectively. Functionally characterized enzymes are labeled at the tree branches. The four AAADs for which their crystal structures were resolved in this study are denoted in bold. The *Eg*PAAS identified and characterized in this study is underlined. * and ** denote two mechnisitic classes of AASs that harbor naturally occurring substitutions at the large-loop catalytic tyrosine or the small-loop catalytic histidine, respectively. (*C*) Taxonomic distribution of plant AAADs across major lineages of green plants. The tree illustrates the phylogenetic relationship among Phytozome V12 species with sequenced genome. The presence of yellow, green, blue or pink circles at the tree branches indicates the presence of one or more AAAD sequences from the bacterial/chlorophyte, basal, TyDC or TDC clades, respectively.

**Fig. 2.**
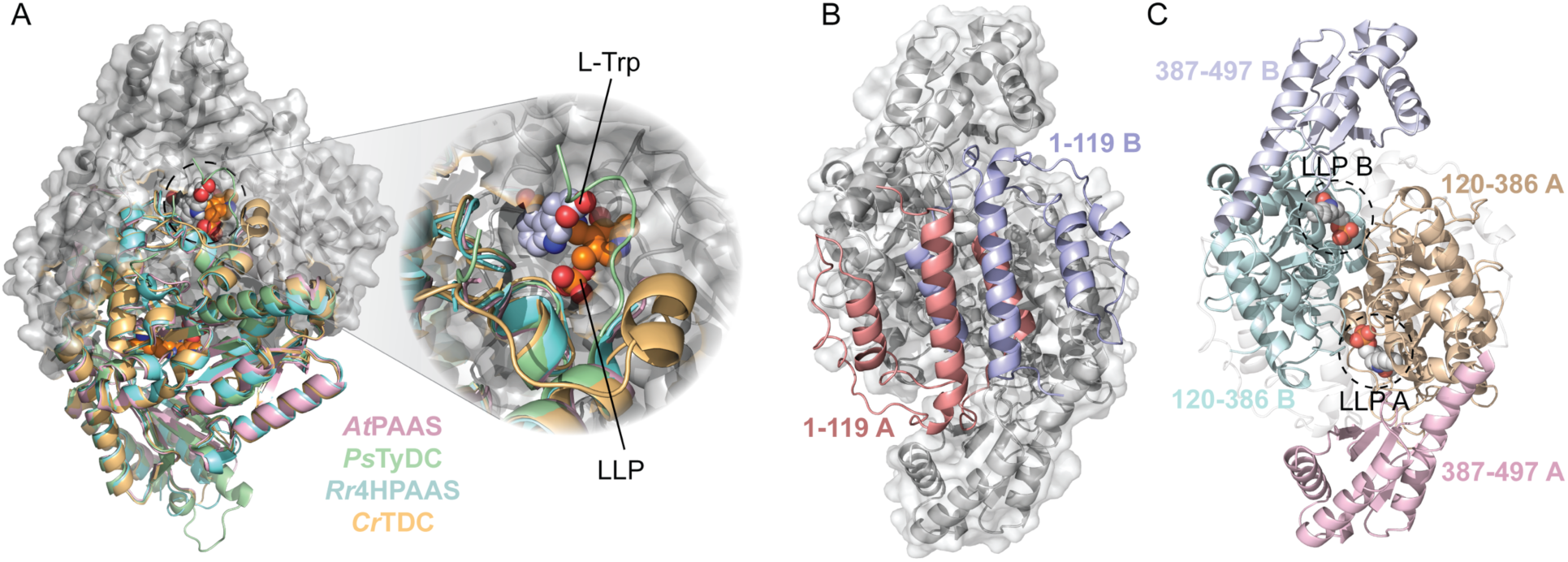
The overall structure of plant AAADs. (*A*) An overlay of the the *Cr*TDC (orange), *Ps*TyDC (green), *At*PAAS (pink), and *Rr*4HPAAS (cyan) structures. All four structures exist as highly similar homodimers, but for visual simplicity, the cartoon structures were only displayed for the bottom monomers. The top monomer of *Cr*TDC is displayed in gray cartoon and surface representation. The dotted circle highlights the *Cr*TDC active site which contains the L-tryptophan substrate and the prosthetic group LLP. (*B*) The *Cr*TDC N-terminal segments from the two monomers, colored in salmon and blue respectively, form the hydrophobic dimer interface. The remainder of the homodimer is displayed in gray. (*C*) The configuration of the *Cr*TDC middle (colored in teal and brown) and C-terminal segments (colored in blue and pink) from the two monomers. The N-terminal segments are displayed in transparent gray cartoons and the prosthetic LLPs circled and displayed as spheres. The models exhibited in (*B*) and (*C*) are in the same orientation, which is rotated 90 degrees around the vertical axis from the view in (*A*).

### Structural features that control substrate selectivity in plant AAADs

The substrate-binding pocket of plant AAADs is principally composed of conserved hydrophobic residues (Trp^92^, Phe^100^, Phe^101^, Pro^102^, Val^122^, Phe^124^, His^318^ and Leu^325^ as in *Cr*TDC) in addition to three variable residues (Ala^103^, Thr^369^ and Gly^370^ as in *Cr*TDC) that display sequence divergence across major AAAD clades (Fig. 3*A, B*). Comparison of the *Cr*TDC and *Ps*TyDC ligand-bound structures rationalizes the role of Gly-versus-Ser variation at position 370 (numbering according to *Cr*TDC) in dictating the size and shape of the substrate-binding pocket to favor indolic versus phenolic amino acid substrate in *Cr*TDC and *Ps*TyDC, respectively (Fig. 3*C*) (11). This observation is consistent with the previous observation that the *Cr*TDC^G370S^ mutant exhibits enhanced affinity for the nonnative substrate L-DOPA as compared to wild-type *Cr*TDC (11). To test how the G370S mutation would impact *Cr*TDC activity *in vivo,* we compared the monoamine product profiles of transgenic yeast expressing wild-type *Cr*TDC or *Cr*TDC^G370S^ measured by liquid chromatography high-resolution accurate-mass mass-spectrometry (LC-HRAM-MS). Compared to the *Cr*TDC-expressing yeast, the *Cr*TDC^G370S^-expressing yeast showed little changes in tryptamine level (Fig. 3*D*) but elevated accumulation of phenylethylamine (Fig. 3*E*), supporting the role of residue at position 370 in gating indolic versus phenolic substrates.

**Fig. 3.**
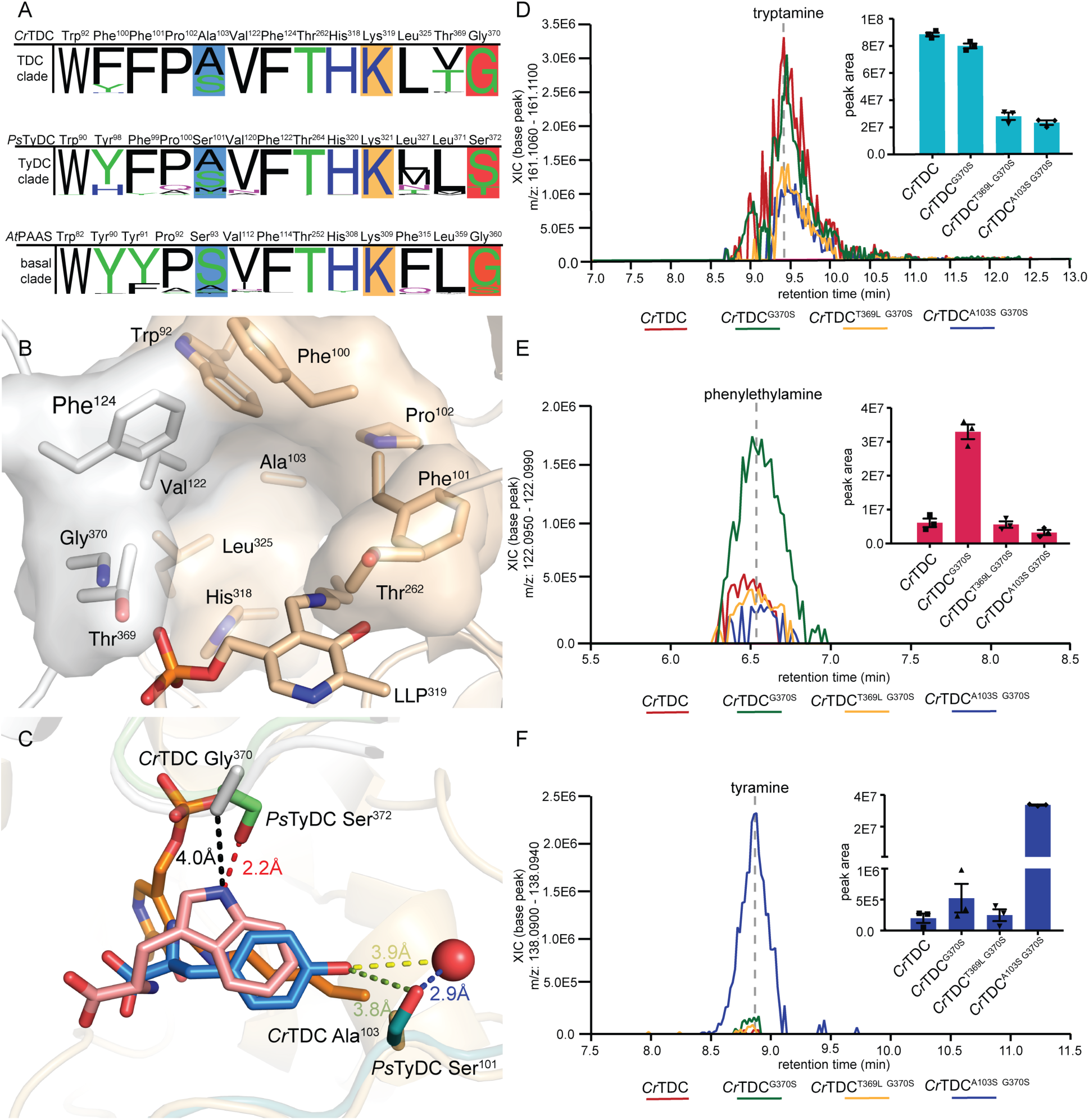
Active site pocket composition and residues that dictate substrate selectivity in plant AAADs. (*A*) Active-site-lining residues from plant AAADs were identified and queried for conservation against all AAAD homologs identified from the 93 Phytozome V12.1 annotated green plant genomes. The height of the residue label displays the relative amino acid frequency, excluding sequence gaps, in the basal, TyDC, and TDC clades. The position of the active site pocket residues from each clade are referenced against the *At*PAAS, *Ps*TyDC and *Cr*TDC respectively. Polar amino acids are colored in green, basic amino acids are colored in blue, acidic amino acids are colored in red, and hydrophobic amino acids are colored in black. Residues highlighted in blue and dark orange boxes denote residues involved in hydroxylated vs. unhydroxylated and phenolic vs. indolic substrate recognition, respectively. The conserved lysine residues, represented as the LLP prosthetic group in several crystal structures, is marked by a light orange box. (*B*) The *Cr*TDC active-site pocket is composed of residues from both chain A and B, colored in beige and white, respectively. The pocket is composed of conserved nonpolar residues (Pro^102^, Val^122^ and Leu^325^), aromatic residues (Trp^92^, Phe^100^, Phe^101^, Phe^124^, His^318^), and a polar residue (Thr^262^). Additionally, the active site contains three variable residues (Ala^103^, Thr^369^ and Gly^370^), which differ across different AAAD clades. (*C*) Superimposition of the substrate-complexed *Cr*TDC and *Ps*TyDC structures. *Cr*TDC Chain A and Chain B are displayed in beige and white, respectively, while relevant portions of the *Ps*TyDC Chain A and Chain B are displayed in green and deep teal, respectively. The L-tryptophan ligand from the *Cr*TDC structure is colored in pink, while the *Ps*TyDC L-tyrosine ligand is colored in blue. The red sphere represents a *Ps*TyDC active-site water likely involved in substrate recognition. Relative *in vivo* activities of L-tryptophan decarboxylase (*D*), L-phenylalanine decarboxylase (*E*), and L-tyrosine decarboxylase (*F*) of wild-type and various mutants of *Cr*TDC were measured in transgenic yeast. The error bars indicate standard error of the mean (SEM) based on biological triplicates.

In the *Cr*TDC structure, the second variable residue Thr^369^ is adjacent to the indolic-selective Gly^370^ residue, which is mostly conserved as a leucine in both the basal clade and the TyDC clade but varies more widely as a threonine, valine, or phenylalanine in the TDC clade (Fig. 3*A*). The variable nature of this residue in the TDC clade suggests a potential role of this residue in substrate selectivity. However, transgenic yeast expressing the *Cr*TDC^T369L G370S^ double mutant showed a general decrease in aromatic monoamine production but no significant difference in relative abundance of each monoamine product compared to yeast strain expressing the *Cr*TDC^G370S^ single mutant (Fig. 3*E-F*). This observation thus did not support a direct correlation of the identity of this second variable residue with substrate selectivity.

The third variable residue at position 103 (numbering according to *Cr*TDC) is represented only as a serine or alanine in the TDC clade but varies as serine, threonine, alanine, methionine, phenylalanine, or cysteine in the basal and TyDC clades (Fig. 3*A*). In the L-tyrosine-bound *Ps*TyDC structure, the *p*-hydroxyl of the tyrosine substrate is coordinated by *Ps*TyDC Ser^101^ (corresponding to *Cr*TDC Ala^103^) together with a nearby water through hydrogen bonding (Fig. 3*C*), suggesting the role of this residue in gating hydroxylated versus unhydroxylated aromatic amino acid substrates. Indeed, transgenic yeast expressing the *Cr*TDC^A101S G370S^ double mutant exhibited an approximately 64-fold increase in tyramine level and a significant decrease in phenylethylamine production as compared to the *Cr*TDC^G370S^-expressing strain (Fig. 3*E-F*). The residue variation at this position in the basal and TyDC clades likely plays a role in discerning various phenolic substrates, namely L-phenylalanine, L-tyrosine and L-DOPA, with varying ring hydroxylation patterns. Serine or alanine substitution at this position in the TDC clade may likewise distinguish 5-hydroxy-L-tryptophan versus L-tryptophan substrate for entering the plant melatonin and indole alkaloid biosynthetic pathways, respectively. Together, these results suggest that the phylogenetically restricted sequence variations at position 103 and 370 (numbering according to *Cr*TDC) were likely selected to control the respective substrate preferences of the TyDC and TDC clades as they diverged from the basal-clade AAAD progenitors.

### The PLP-dependent catalytic center of plant AAADs

PLP, the active form of vitamin B6, is a versatile coenzyme employed by approximately 4% of all known enzymes to catalyze a diverse array of biochemical reactions. The active site of canonical AAADs constrains the reactivity of PLP to specifically catalyze decarboxylation of the α-carbon-carboxyl group of aromatic L-amino acids (12). The active site of plant AAADs, as represented in the *Cr*TDC structure, is located at the dimer interface, and features the characteristic prosthetic group LLP in the form of an internal aldimine at resting state (Fig. S7). The phosphate moiety of LLP is coordinated by Thr^167^, Ser^168^ and Thr^369^, while the pyridine-ring amine forms a salt bridge with the side-chain carboxyl of Asp^287^, supporting its conserved role in stabilizing the carbanionic intermediate of the PLP external aldimine (Fig. S7) (22). Evident from the L-tryptophan-bound *Cr*TDC structure, the aromatic L-amino acid substrate is oriented in the active site to present its labile Cα-carboxyl bond perpendicular to the pyridine ring of the internal aldimine LLP (Fig. S8) as predicted by Dunathan’s hypothesis (12). Upon substrate binding, transaldimination then occurs that conjugates the α-amino group of the substrate to PLP through a Schiff base to yield the external aldimine (Fig. S9), as likely captured by one of the active sites of the L-tyrosine-bound *Ps*TyDC structure (Fig. S10). The resulting external aldimine subsequently loses the α-carboxyl group as CO_2_ to generate a quinonoid intermediate (Fig. 4*A*, reaction step 1). Through an as-yet-unknown mechanism, the nucleophilic carbanion of the quinonoid intermediate is protonated to yield the monoamine product and a regenerated LLP, ready for subsequent rounds of catalysis (Fig. 4*A*, reaction steps 2-3) (13). Despite the availability of several animal DDC crystal structures, several aspects of the full catalytic cycle of the type II PLP decarboxylases remain speculative. Comparative structural analysis of our four plant AAAD structures sheds new light on the mechanistic basis for the canonical decarboxylation activity as well as the evolutionarily new decarboxylation-dependent oxidative deamination activity, which are detailed below.

**Fig. 4.**
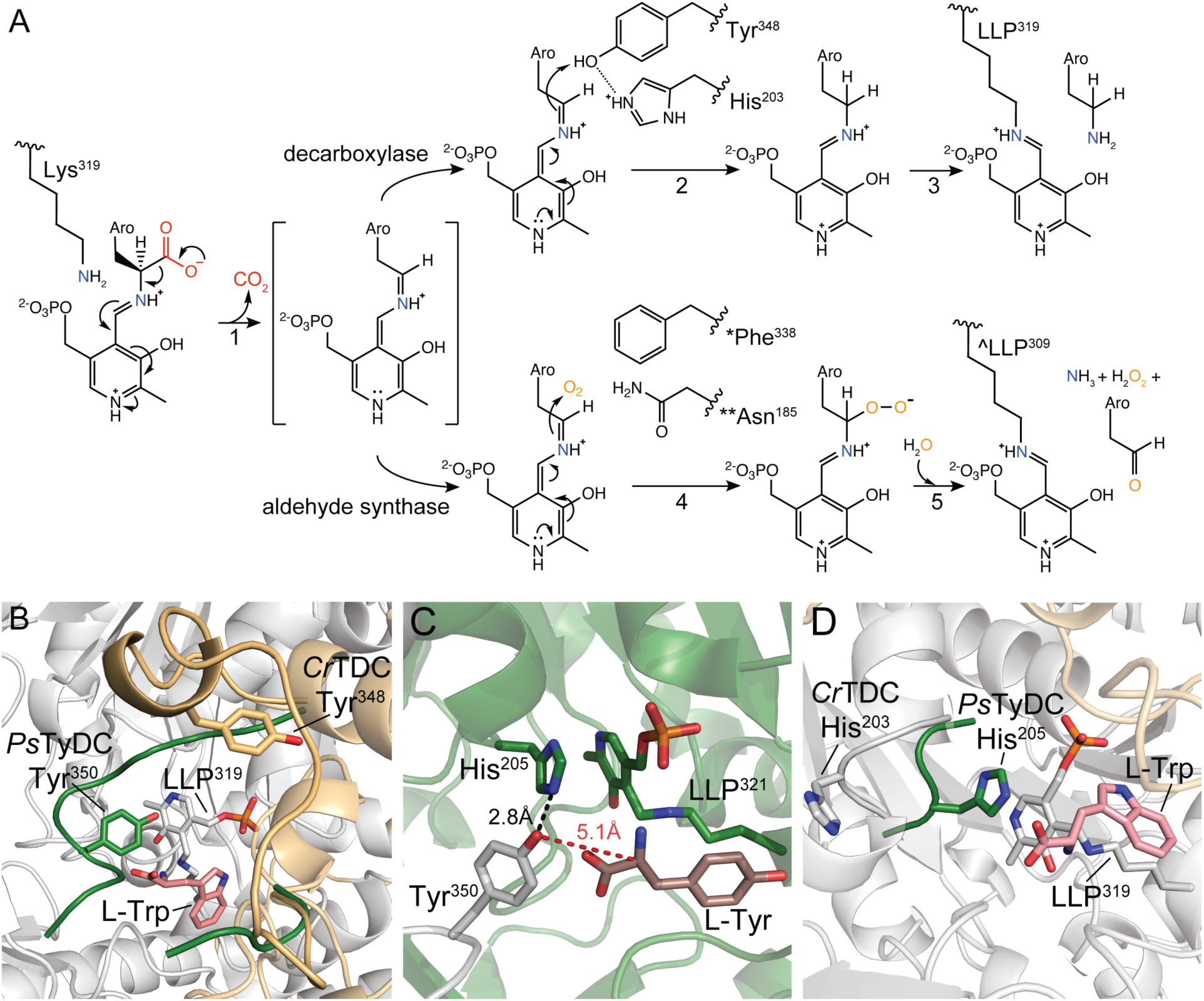
Catalytic mechanisms and conformational changes of plant AAAD proteins. (*A*) The proposed alternative PLP-mediated catalytic mechanisms for the canonical decarboxylase and derived aldehyde synthase in plant AAAD proteins. After transaldimation of the *Cr*TDC internal aldimine to release the active-site Lys^319^, the PLP-amino-acid external aldimine loses the α-carboxyl group as CO_2_ to generate a quinonoid intermediate stabilized by the delocalization of the paired electrons (**1**). In a canonical decarboxylase (e.g., *Cr*TDC), the carbanion at Cα is protonated by the acidic *p*-hydroxyl of Tyr^348-B^ located on the large loop, which is facilitated by its neighboring His^203-A^ located on the small loop (**2**). The *Cr*TDC LLP^319^ internal aldimine is regenerated, accompanied by the release of the monoamine product (**3**). In an evolutionarily new aldehyde synthase, the Cα protonation step essential for the canonical decarboxylase activity is impaired, when the large-loop catalytic tyrosine is mutated to phenylalanine (* as in *At*PAAS), or when the small-loop catalytic histidine is mutated to asparagine (** as in *Eg*PAAS), allowing the Cα carbanion to attack a molecular oxygen to produce a peroxide intermediate (**4**). This peroxide intermediate decomposes to give aromatic acetaldehyde, ammonia, and hydrogen peroxide products and regenerate the LLP internal aldimine (^ as in *At*PAAS) (**5**). Aro represents the aromatic moiety of an aromatic L-amino acid. (*B*) An overlay of the *Ps*TyDC structure with its large loop in a closed conformation (green) upon the *Cr*TDC structure with its large loop in an open conformation (beige). (*C*) The closed conformation of *Ps*TyDC active site displaying the catalytic machinery in a configuration ready to engage catalysis. Chain A is colored in white, chain B is colored in green, and the L-tyrosine substrate is displayed in dark pink. (*D*) Open and closed small loop conformations observed in the *Cr*TDC and *Ps*TyDC structures. The *Ps*TyDC small loop with the catalytic histidine is in a closed conformation (green), while the *Cr*TDC structure exhibits its small-loop histidine in an open conformation (white).

### Structural features of two catalytic loops

Previous structural studies of the mammalian AAADs identified a large loop that harbors a highly conserved tyrosine residue essential for catalysis (14). However, this loop is absent from the electron density map of all the previously resolved animal DDC structures. We provide the first direct electron-density support for this large loop that adopts different conformations relative to the active site in different plant AAAD crystallographic datasets (Fig. 4*B* and Supplementary Note 1). In the *Cr*TDC structure, the large loop (*Cr*TDC^342-361^) of monomer A adopts an open conformation, revealing a solvent-exposed active site. Conversely, in the *Ps*TyDC structure, a crankshaft rotation puts the large loop over the bound L-tyrosine substrate to seal the active-site pocket. In this closed conformation, the *p*-hydroxyl of the catalytic tyrosine located on the large loop (Tyr^350-A^) is in close vicinity with the Cα of the L-tyrosine substrate and LLP (Fig. 4*C*). These structural observations suggest that Tyr^350^ (as in *Ps*TyDC) serves as the catalytic acid that protonates the nucleophilic carbanion of the quinonoid intermediate at the substrate Cα position, which is preceded by an open-to-closed conformational change by the large loop.

Comparative structural analysis of our four plant AAAD structures identified another small loop (*Cr*TDC^200-205-B^), which harbors a conserved histidine residue (His^203-B^ as in *Cr*TDC) and cooperatively interacts with the large loop from the opposite monomer. For instance, the small loop in the *Cr*TDC structure is rotated outward from the active-site LLP (Fig. 4*D*). Conversely, in the *At*PAAS, *Ps*TyDC and *Rr*4HPAAS structures, the small loop adopts a closed conformation with its histidine imidazole forming pi stacking with the LLP pyridine ring. Such pi stacking between PLP and an active-site aromatic residue has been observed previously as a common feature within AAADs as well as the broader α-aspartate aminotransferase superfamily (15–17). Moreover, in the *Ps*TyDC structure, the τ-nitrogen of the small-loop histidine is in hydrogen-bonding distance with the *p*-hydroxyl of the large-loop catalytic tyrosine (Fig. 4*C*), suggesting a potential catalytic role of this histidine. Previous studies of the pH dependence of AAADs suggest that this small-loop histidine can not function directly as the catalytic acid to protonate the carbanionic quinonoid intermediate (18–20). Based on our structural observations, we propose a catalytic mechanism where the small-loop histidine (*Cr*TDC His^203-B^) facilitates the proton transfer from the *p*-hydroxyl of the large-loop tyrosine (*Cr*TDC Tyr^348-A^) to the quinonoid intermediate (Fig. 4*A*, reaction steps 2) by serving as a direct or indirect proton source for reprotonation of the Tyr^348-A^ *p*-hydroxyl. Mechanistically similar Tyr-His side-chain interaction that facilitates protonation of a carbanionic intermediate has also been recently proposed for the *C. roseus* heteroyohimbine synthase, which is a medium-chain dehydrogenase/reductase-family enzyme (21).

### Molecular dynamics simulations reveal the dynamic nature of the two catalytic loops

Considering the various alternative conformations observed in our plant AAAD structures, we sought to examine the flexibility and cooperativity of the two loops that harbor the key catalytic residues (*Cr*TDC Tyr^348-A^ and His^203-B^) by MD simulation. We began with 36 sets of 100-ns simulations on six *Cr*TDC systems with LLP and the substrate L-tryptophan in different protonation states (Fig. S11 and Supplementary Note 2). These simulations revealed considerable flexibility of both loops, with one simulation capturing a dramatic closing motion of the open large loop. Upon extending this simulation to 550 ns (Fig. S12 and Video S1), the large loop was found to reach a semi-closed state characterized by a minimal C_α_ RMSD of 4.3 Å with respect to the modeled *Cr*TDC closed-state structure (Supplementary Note 3). The catalytic Tyr^348-A^ was found to form stacking interactions with His^203-B^, with a minimal distance of approximately 2 Å between the two residues (Fig. S13). Interestingly, a large-loop helix (*Cr*TDC^346-350A^) that unfolded at the beginning of this simulation appeared to ‘unlock’ the large loop from its open state. To further examine the correlation between the secondary structure of this helix and the large-loop conformation, we artificially unfolded this short helix and initiated 72 sets of 50-ns simulations from the resulting structure. Similar conformations to that identified in Fig. S11 were observed in all six systems (Fig. S14*C*) with minimal C_α_ RMSD ranging from ∼5 to 8 Å with respect to the modeled *Cr*TDC closed-state structure. These results suggest that the initial closing motion of the large loop is independent of the coenzyme and substrate protonation states and that the unfolding of the aforementioned helix can significantly accelerate such motion. This is further substantiated by three sets of 600-ns simulations of *Cr*TDC in an apo state with neither PLP nor L-tryptophan present (Fig. S15 and Supplementary Note 2). It is noted, however, that the fully closed state of *Cr*TDC was not achieved in our sub-microsecond simulations. For instance, in the trajectory shown in Fig. S11 and Video S1, L-tryptophan left the active site at around t=526 ns, shortly after which the simulation was terminated. Overall, while the transition from the semi-closed to the fully closed state can be expected to occur beyond the sub-microsecond timescale, the MD results support our hypothesis that Tyr^348-A^ can form close contact with His^203-B^, readying the latter residue to stabilize pyridine ring resonance and direct proton transfer from the former residue to the carbanionic quinonoid intermediate.

### Convergent evolution of two mechanistic classes of AASs

Our new insights into the structural and dynamic basis for the canonical AAAD catalytic mechanism predicts that mutations to the large-loop tyrosine or the small-loop histidine would likely derail the canonical AAAD catalytic cycle and potentially yield alternative reaction outcomes. Indeed, the canonical large-loop tyrosine is substituted by phenylalanine in *At*PAAS and *Rr*4HPAAS. We propose that absence of the proton-donating *p*-hydroxyl of the catalytic tyrosine enlarges the active-site cavity to permit molecular oxygen to enter the active site and occupy where the tyrosine *p*-hydroxyl would be for proton transfer (Fig. 4*A*, reaction steps 4-5, *). Instead of protonation, the nucleophilic carbanion of the quinonoid intermediate attacks molecular oxygen to generate a peroxide intermediate, which subsequently decomposes to yield ammonia, hydrogen peroxide and the aromatic acetaldehyde product.

The rapid expansion of genomic resources together with our increasing knowledge about the structure-function relationships of disparate enzyme families enables database mining for potentially neofunctionalized enzymes in plant specialized metabolism. Indeed, the identification of the Tyr-to-Phe substitution responsible for the AAS activity in plants (22) served as a “molecular fingerprint” for the functional prediction of unconventional AASs among legume serine decarboxylase-like enzymes (23). Bearing in mind the newly identified functional residues that control substrate selectivity and catalytic mechanism, we queried all plant AAAD homologs identifiable by BLAST searches within NCBI and the 1KP databases (24). Interestingly, this search identified a number of TyDC-clade AAAD homologs from the Myrtaceae and Papaveraceae plant families that harbor unusual substitutions at the small-loop histidine (Fig. S16). Several Myrtaceae plants contain a group of closely related and likely orthologous AAAD proteins with a His-to-Asn substitution at the small-loop histidine. In Papaveraceae, on the other hand, we noticed a His-to-Leu substitution in the previously reported *P. somniferum* tyrosine decarboxylase 1 (*Ps*TyDC1) (8) and a His-to-Tyr substitution in another *Papaver bracteatum* AAAD homolog. While the small-loop histidine is typically conserved among type II PLP decarboxylases and the broader type I aspartate aminotransferases, the same His-to-Asn substitution was previously observed in select insect AAAD proteins involved in 3,4-dihydroxyphenylacetaldehyde production necessary for insect soft cuticle formation (25, 26). These sequence observations together with the proposed catalytic role of the small-loop histidine implicate that AAADs with substitutions at this highly conserved residue may contain alternative enzymatic functions.

To test this hypothesis, we first generated transgenic yeast strains expressing either wild-type *Ps*TyDC, *Ps*TyDC^Y350F^, or *Ps*TyDC^H205N^, respectively, and assessed their metabolic profiles by LC-HRAM-MS (Fig. 5*A-B*). The *Ps*TyDC-expressing yeast exclusively produces tyramine, whereas the *Ps*TyDC^Y350F^-expressing yeast exclusively produces tyrosol (reduced from 4HPAA by yeast endogenous metabolism). This result suggests that mutating the large-loop tyrosine to phenylalanine in a canonical TyDC sequence background is necessary and sufficient to convert it to an AAS. Interestingly, the *Ps*TyDC^H205N^-expressing yeast produces both tyramine and tyrosol, suggesting that mutating the small-loop histidine to asparagine alone in *Ps*TyDC turned it into an AAAD-AAS bifunctional enzyme. *In vitro* enzyme assays using recombinant wild-type *Ps*TyDC, *Ps*TyDC^Y350F^, or *Ps*TyDC^H205N^ against L-tyrosine as substrate yielded similar results that corroborated the observations made in transgenic yeast (Fig. 5*C*). Collectively, these results reinforce the essential role of the large-loop catalytic tyrosine for the canonical AAAD activity, and support the proposed assisting role of the small-loop histidine in quinonoid intermediate protonation for the canonical AAAD activity. The full AAAD catalytic cycle could still proceed, albeit with significantly reduced efficiency, when the small-loop histidine is mutated to asparagine, whereas a significant fraction of the carbanionic quinonoid intermediate undergoes the alternative oxidative deamination chemistry similarly as proposed for *At*PAAS (Fig. 4*A*, reaction steps 4-5, **).

**Fig. 5.**
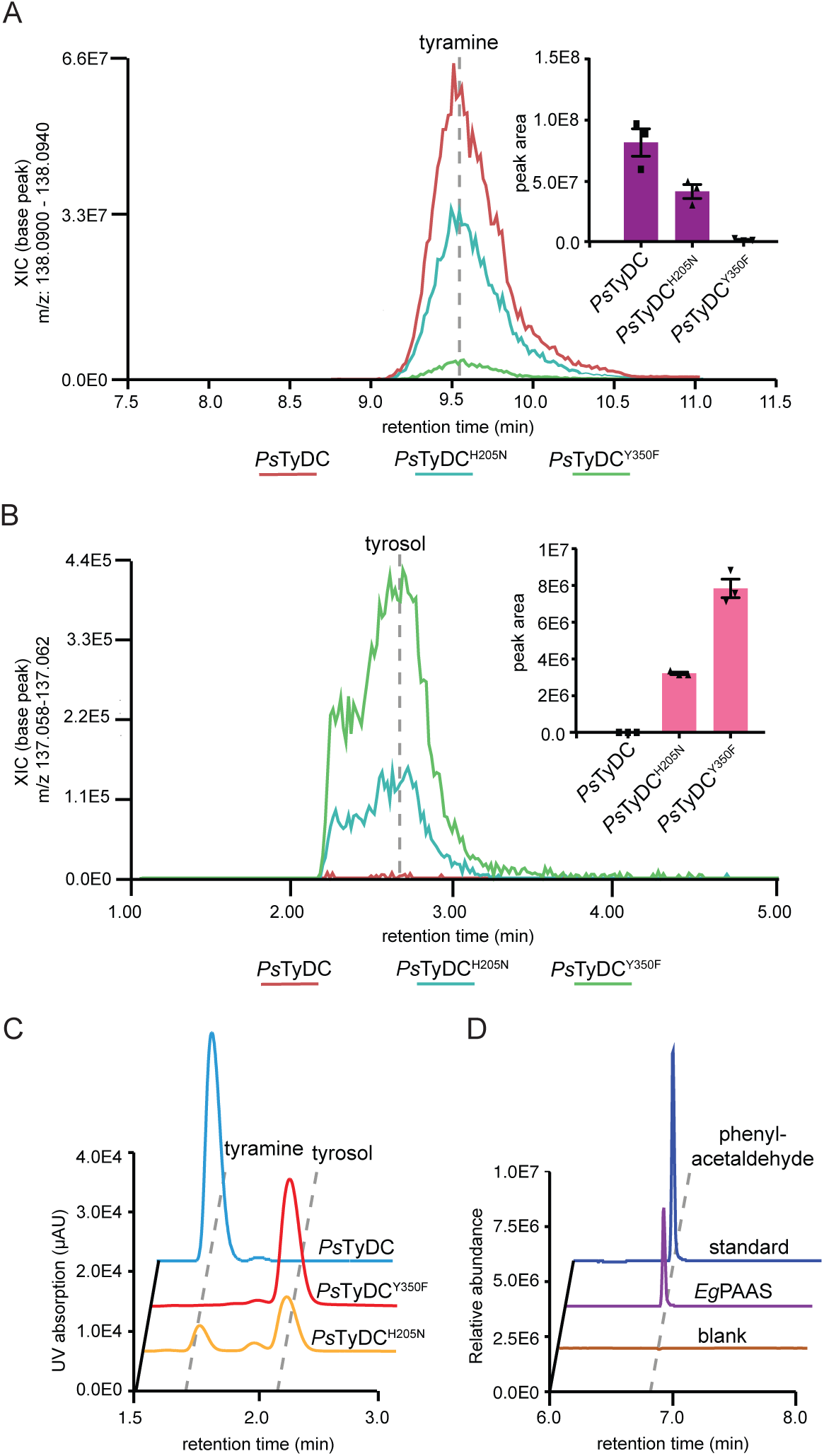
Two alternative molecular strategies to arrive at aldehyde synthase chemistry from a canonical AAAD progenitor. Transgenic yeast strains expressing *Ps*TyDC^H205N^ and *Ps*TyDC^Y350F^ display reduced tyramine production (*A*) but elevated levels of tyrosol (*B*) (reduced from 4HPAA by yeast endogenous metabolism) in comparison to transgenic yeast expressing wild-type *Ps*TyDC. Cultures were grown and metabolically profiled in triplicate, and the error bars in the bar graphs indicate standard error of the mean (SEM). (*C*) LC-UV chromatograms showing the relative decarboxylation and aldehyde synthase products produced by purified recombinant *Ps*TyDC, *Ps*TyDC^Y350F^or *Ps*TyDC^H205N^ enzymes when incubated with 0.5 mM L-tyrosine for 5, 25, and 50 minutes, respectively. After enzymatic reaction, the 4HPAA aldehyde product was chemically reduced by sodium borohydride to tyrosol prior to detection. (*D*) Phenylacetaldehyde formation from the incubation of purified recombinant *Eg*PAAS with L-phenylalanine measured by GC-MS.

To assess the function of plant AAAD homologs that naturally contain substitutions at the small-loop histidine, we cloned one of such genes from *Eucalyptus grandis* (*EgPAAS*, Fig. 1*B*), starting from total mRNA extracted from the host plant. *In vitro* enzyme assays conducted using recombinant *Eg*PAAS against a panel of aromatic L-amino acids demonstrated exclusive AAS activity with an apparent substrate preference towards L-phenylalanine (Fig. 5*D* and Fig. S17). It is worth noting that phenylacetaldehyde has been previously identified as a major fragrance compound present in the essential oil and honey of numerous Myrtaceae plants (27–29), which may be attributed to the activity of *Eg*PAAS. Unlike the *Ps*TyDC^H205N^ mutant, *Eg*PAAS shows no detectable ancestral AAAD activity *in vitro*, suggesting that in addition to the small-loop His-to-Asn substitution, other adaptive mutations must have contributed to the specific PAAS activity. While the AAS activity of *Eg*PAAS was confirmed in transgenic yeast (Fig. S18), Myrtaceae AAADs from *Medinilla magnifica* and *Lagerstroemia indica* (*Mm*AAS and *Li*AAS), which also contain the unusual small-loop Hist-to-Asn substitution (Fig. S16), displayed little to no enzymatic activity in this system. Despite our interest in the other two AAAD sequences from Papaveraceae plants harboring alternative substitutions at the small-loop histidine, they could not be directly cloned from their respective host plants, raising the possibility that these genes could potentially be derived from sequencing or assembly errors. Thus, we did not pursue functional characterization of these genes further.

### Metabolic engineering of a shortened BIA biosynthetic pathway in yeast with 4HPAAS

To examine the role of adaptive functional evolution of a single AAAD protein in the larger context of specialized metabolic pathway evolution, we attempted a metabolic engineering exercise to build a shortened BIA biosynthetic pathway in yeast *S. cerevisiae*. In the proposed plant BIA pathway, the key intermediate 4HPAA is reportedly derived from L-tyrosine through two consecutive enzymatic steps catalyzed by L-tyrosine aminotransferase (TAT) and an unidentified 4-hydroxyphenylpyruvate decarboxylase (4HPPDC) (30) (Fig. 6*A*). However, in the *R. rosea* salidroside biosynthetic pathway, 4HPAA is directly converted from L-tyrosine by a single *Rr*4HPAAS enzyme (10) (Fig. 6*A*). We therefore reasoned that 4HPAA activity could be utilized to reroute the 4HPAA branch of the BIA pathway. In our metabolic engineering scheme, either *Rr*4HPAAS or the *Ps*TyDC^Y350F^ mutant was used as the 4HPAAS enzyme, while wild-type *Ps*TyDC was used as a control. Moreover, *Pseudomonas putida* DDC (*Pp*DDC) and *Beta vulgaris* L-tyrosine hydroxylase (*Bv*TyH) were used to generate the other key intermediate dopamine from L-tyrosine. Lastly, *P. somniferum* NCS (*Ps*NCS) was added to stereoselectively condense 4HPAA and dopamine to form (S)-norcoclaurine.

**Fig. 6.**
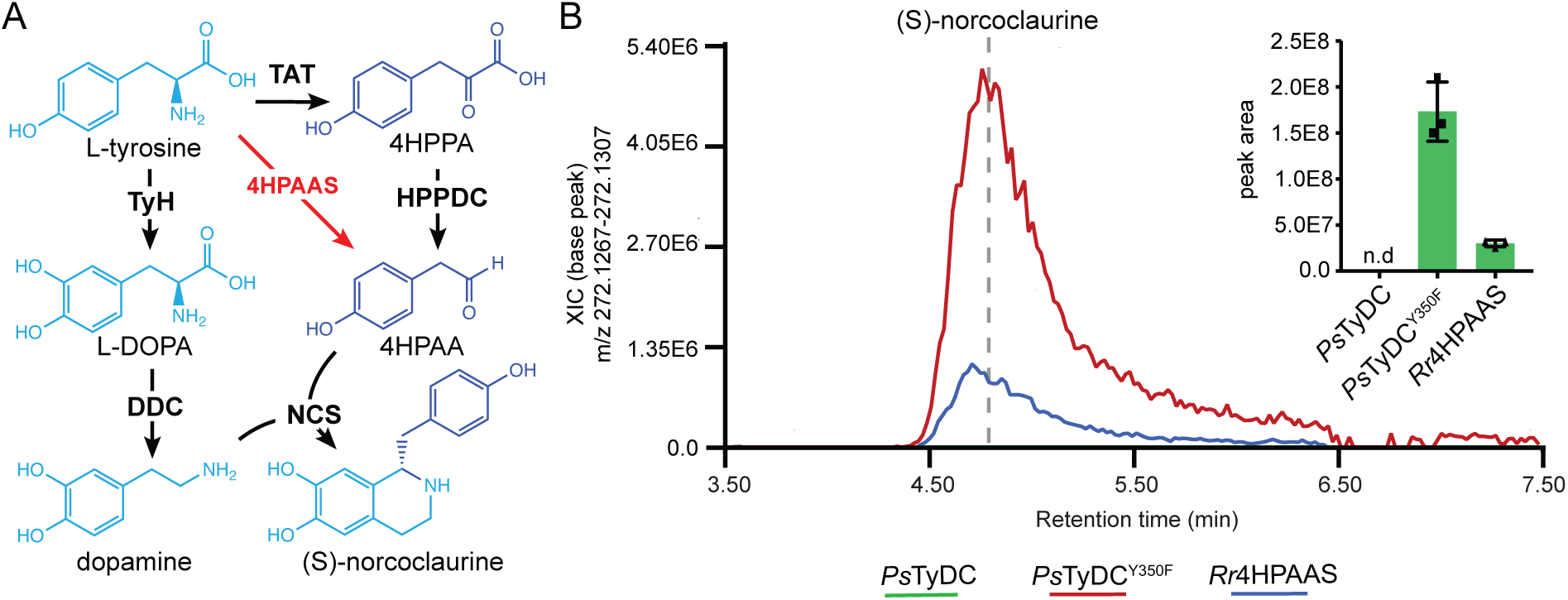
The utility of 4HPAAS in metabolic engineering of a shortened BIA pathway for (S)-norcoclaurine production in yeast. (*A*) Canonically proposed (S)-norcoclaurine biosynthetic pathway (black arrows) rerouted by the use of an 4HPAAS (red arrows). 4HPPA, 4-hydroxyphenylacetaldehyde. (*B*) Engineering of (S)-norcoclaurine production in yeast using two AAAD proteins with the 4HPAAS activity. All multigene vectors used to transform yeast contain *BvTyH*, *PpDDC*, and *PsNCS* in addition to either *PsTyDC*, *PsTyDC*^Y350F^ or *Rr4HPAAS*. Cultures were grown and measured in triplicate, and the error bars in the bar graph indicate standard error of the mean (SEM). n.d., not detected.

Coexpression of *Ps*TyDC^Y350F^ with *Pp*DDC, *Bv*TyH and *Ps*NCS resulted in the highest accumulation of (S)-norcoclaurine among various engineered transgenic yeast strains, whereas replacing *Ps*TyDC^Y350F^ with *Rr*4HPAAS gave approximately 7-fold less (S)-norcoclaurine production (Fig. 6*B*). Control experiment using wild-type *Ps*TyDC produced tyramine but not (S)-norcoclaurine as expected (Fig. 6*B* and Fig. S19). This metabolic engineering exercise illustrates that, when provided with the right metabolic network context, the birth of an evolutionarily new AAS activity (achievable through a single mutation) can lead to rewiring of the ancestral specialized metabolic pathway. Mechanistic understanding of the structure-function relationship within the versatile plant AAAD family adds to the toolset for metabolic engineering of high-value aromatic-amino-acid-derived natural products in heterologous hosts.

## Discussion

Selective pressures unique to plants’ ecological niches have shaped the evolutionary trajectories of the rapidly expanding specialized metabolic enzyme families in a lineage-specific manner. The AAAD family is an ancient PLP-dependent enzyme family ubiquitously present in all domains of life. While AAAD proteins mostly retain highly conserved primary metabolic functions in animals, e.g. monoamine neurotransmitter synthesis, they have undergone extensive radiation and functional diversification during land plant evolution. We show that the major monophyletic TDC and TyDC clades of plant AAADs emerged from the basal clade, each acquiring divergent active-site structural features linked to substrate specificity. Moreover, the ancestral AAAD catalytic machinery has also been modified repeatedly through two types of mechanistic mutations to converge on the evolutionarily new AAS activity in numerous AAAD paralogs in plants and insects. The rich evolutionary history of the AAAD family therefore affords a prism for understanding how plants capitalize their ability to biosynthesize proteinogenic aromatic L-amino acids and use them as precursors for further developing taxonomically restricted specialized metabolic pathways (31). Some of these downstream pathways, such as the tryptophan-derived MIA pathway in plants under the order of Gentianales and the tyrosine-derived BIA pathway in plants under the order of Ranunculales, give rise to major classes of bioactive plant natural products critical not only for host fitness but also as a source for important human medicines.

PLP in isolation can catalyze almost all of the reactions known for PLP-dependent enzymes, including transamination, decarboxylation, 0- and y-elimination, racemization and aldol cleavage (32). In the context of PLP-dependent enzymes, PLP reactivity is confined by its encompassing active site to elicit specific catalytic outcome with defined substrates (32). Evolutionary analysis suggests that major functional classes of PLP-dependent enzymes likely established first, followed by divergence in substrate selectivity (32, 33). Four evolutionarily distinct groups of PLP-dependent decarboxylases exist in nature (33). The largest and most diverse group (Group II) consists of AAADs, histidine decarboxylases, glutamate decarboxylases, aspartate decarboxylases, serine decarboxylases, and cysteine sulfinic acid decarboxylases, implicating a deep divergence of amino acid substrate selectivity among Group II enzymes. Within plant AAADs, the substrate selectivity has continued to evolve to arrive at extant enzymes with exquisite substrate selectivity towards various proteinogenic and non-proteinogenic aromatic L-amino acids. This is largely achieved by fixing specific mutations at the substrate-binding pocket as revealed in this study.

Oxygenation is one of the most common chemical reactions in plant specialized metabolism. The triplet ground electron state of molecular oxygen necessitates a cofactor to facilitate single-electron transfer, evident from several major oxygenase families involved in plant specialized metabolism, including cytochrome P450 monooxygenases (P450s), iron/2-oxoglutarate-dependent oxygenases (Fe/2OGs), and flavin-dependent monooxygenases (FMOs) (34). AASs represent a class of plant oxygenases that exploit a carbanionic intermediate to facilitate oxygenation (Fig. 4*A*, reaction steps 4-5). This catalytic mechanism is reminiscent of the paracatalytic photorespiratory reaction catalyzed by ribulose-1,5-bisphosphate carboxylase/oxygenase (RuBisCO), where molecular oxygen instead of CO_2_ enters the RuBisCO active site and reacts with a ribulose-1,5-bisphosphate-derived carbanionic enediolate intermediate to form a peroxide intermediate that subsequently decomposes to yield 3-phosphoglycerate and 2-phosphoglycerate (35). Carbanion-stabilizing enzymes, including PLP-dependent enzymes DDC, glutamate decarboxylase and ornithine decarboxylase, are also capable of catalyzing similar paracatalytic oxygenation reactions (35–38). Whereas oxygenation reactions observed in these carbanion-stabilizing enzymes are mostly deemed as paracatalytic activities due to the intrinsic reactivity of the carbanion intermediate, the plant AASs evolved by harnessing such latent paracatalytic activity for dedicated production of aromatic acetaldehydes and their derived secondary metabolites (10, 39–42). We show that, by understanding the structural and mechanistic basis for convergent evolution of two mechanistic classes of AAS within the plant AAAD family, *Ps*TyDC could be converted to a specific 4HPAAS with a single large-loop Tyr-to-Phe mutation, or an AAAD-AAS bifunctional enzyme by the small-loop His-to-Asn mutation. Several aspects of the neofunctionalization of the AAS activities remain unclear and are topics for future investigation. For example, the emergence of AAS activity results in the stoichiometric production of ammonium, hydrogen peroxide, and reactive aldehydes, which may require additional adaptive metabolic and cellular processes for taming their reactivity/toxicity in AAS-expressing cells. While the TDC and TyDC clades appear to clearly function in specialized metabolic pathways, the biochemical functions and physiological roles of the basal clade in plants remain less studied. Furthermore, the exact chemical mechanism underlying decomposition of the peroxide intermediate that yields aromatic acetaldehyde, ammonium, hydrogen peroxide and the regenerated active-site LLP is yet to be defined.

Co-opting progenitor enzymes to synthesize novel and adaptive metabolites is a universal mechanism underscoring metabolic evolution (43). Most specialized metabolic enzymes present in extant plants evolved through the recruitment of malleable ancestral enzyme folds followed by neofunctionalization of substrate specificity, product diversity, or, in much rarer cases, alternative catalytic mechanisms (44–47). The plant AAAD family illustrates all these evolutionary mechanisms. Applying the learnt knowledge about AAAD evolution has further enabled metabolic engineering of a shortened BIA pathway to produce (S)-norcoclaurine in yeast, using a natural or an evolved 4HPAAS. The use of an insect L-DOPA-specific AAS in engineering of tetrahydropapaveroline biosynthesis in *E. coli* was recently reported (48), highlighting another example of utilizing AAS in metabolic engineering. It is noted that the free aldehydes produced by AASs readily react with amino acids or other free amines to produce iminium-conjugation products via non-enzymatic aldehyde-amine condensation chemistry as seen in *E. coli* (48) or in plant betacyanin and betaxanthin biosynthesis (49). The use of *Ps*NCS that catalyzes the enantioselective Pictet-Spengler condensation of 4HPAA and dopamine (50) therefore helps to channel the reactive 4HPAA towards (S)-norcoclaurine production in engineered yeast. The successful engineering of a shortened (S)-norcoclaurine biosynthetic pathway using 4HPAAS also hints an alternative hypothesis to the currently unresolved plant BIA pathway regarding the origin of 4HPAA (30).

## Materials and Methods

### Reagents

L-tryptophan, tryptamine, L-tyrosine, tyramine, tyrosol, L-phenylalanine, phenylethylamine, phenylacetaldehyde, phenylethyl alcohol, tyrosol, L-3,4-dihydroxyphenylalanine, dopamine, (S)-norcoclaurine, PLP, and sodium borohydride were purchased from Sigma-Aldrich. 4-hydroxyphenylacetaldehyde was purchased from Santa Cruz Biotechnology.

### Multiple sequence alignment and phylogenetic analysis

ClustalW2 was used to generate the protein multiple sequence alignments with default settings (51). The phylogeny shown in Fig. S1 and Fig. S3 were inferred using the Maximum-Likelihood method. The bootstrap consensus unrooted trees were inferred from 500 replicates to represent the phylogeny of the analyzed enzyme families. The phylogenetic analysis encompases AAAD homolog sequences from all sequenced plant genomes available at Phytozome V12 as well as previously characterized AAAD proteins from plants and select bacteria, chordata, and insect sequences (52). All phylogenetic analyses were conducted in MEGA7 (53). ESPript 3.0 was used to display the multiple sequence alignment (54). Conservation of the active-site residues in various AAAD clades was displayed using WebLogo (55).

### Plant materials

*E. grandis* seeds were purchased from Horizon Herbs. Seeds were stratified at 4 °C for three days, and germinated in potting soil. Plants were grown under a 16-h-light/8-h-dark photoperiod at 23 °C in a local greenhouse.

### Molecular cloning

Leaf tissue of seventy-day-old *E. grandis* plants was harvested for total RNA extraction using the Qiagen’s RNeasy Mini Kit (Qiagen). Total RNAs for *A. thaliana*, *P. somniferum, C. roseus* and *R. rosea* were extracted as previously described (56, 57). First-strand cDNAs were synthesized by RT-PCR using total RNA as template. The coding sequences (CDS) of candidate genes were amplified from cDNAs by PCR using gene-specific primers (Table S3). Gibson assembly was used to ligate the *CrTDC*, *PsTyDC*, *AtPAAS* and *Rr4HPAA*S PCR amplicons into pHis8-4, a bacterial expression vector containing an N-terminal 8xHis tag followed by a tobacco etch virus (TEV) cleavage site for recombinant protein production. *EgPAAS* was alternatively cloned though Gibson assembly into pTYB12, a commercially available N-terminal intein/chitin domain fusion vector designed for affinity chromatography purification.

### Recombinant protein production and purification

BL21(DE3) *E. coli* containing the pHis8-4 or pTYB12-based constructs were grown in terrific broth (TB) at 37 °C to OD^600^ of 0.9 and induced with 0.15 mM isopropyl-β-D-thiogalactoside (IPTG). The cultures were cooled to 18 °C and shaken for an additional 20 hours. Cells were harvested by centrifugation, washed with phosphate-buffered saline (PBS) (137 mM NaCl, 2.7 mM KCl, 10 mM Na_2_HPO_4_ and 1.8 mM KH_2_PO_4_), resuspended in 150 mL of lysis buffer (50 mM Tris pH 8.0, 0.5 M NaCl, 20 mM imidazole, and 0.5 mM dithiothreitol (DTT)), and lysed with five passes through an M-110L microfluidizer (Microfluidics). The resulting crude protein lysate from the *Cr*TDC, *Ps*TyDC, *At*PAAS, and *Rr*4HPAAS cultures were clarified by centrifugation prior to Qiagen Ni-NTA gravity flow chromatographic purification. After loading the clarified lysate, His-tagged recombinant protein-bound Ni-NTA resin was washed with 20 column volumes of lysis buffer, and eluted with 1 column volume of elution buffer (50 mM Tris pH 8.0, 0.5 M NaCl, 250 mM imidazole and 0.5mM DTT). 1 mg of His-tagged TEV protease was added to the eluted protein, followed by dialysis at 4 °C for 16 h in dialysis buffer (50 mM Tris pH 8.0, 0.1 M NaCl, 20 mM imidazole and 2 mM DTT). After dialysis, the protein solutions were passed through Ni-NTA resin to remove uncleaved protein and the His-tagged TEV. The *Eg*PAAS enzyme was insoluble when expressed as an N-terminal polyhistidine-tagged protein, and was therefore expressed as a fusion protein with the Intein/Chitin Binding Protein using the pTYB12 vector. *Eg*PAAS-expressing *E. coli* cell pellets were homogenized in an imidazole-free lysis buffer. The resulting crude protein lysate was then applied to a column packed with chitin beads, washed with 1 L of buffer and subsequently hydrolyzed under reducing conditions as per the manufacturer’s instructions. Recombinant proteins were further purified by gel filtration on a fast protein liquid chromatography (FPLC) system (GE Healthcare Life Sciences). The principle peaks were collected, verified for molecular weight by SDS-PAGE, stored in storage buffer (20 mM Tris pH 8.0, 25 mM NaCl, 200 µM PLP and 0.5 mM DTT) at a protein concentration of 10 mg/mL and flash frozen for subsequent investigation. Despite the use of a solubilizing domain from the pTYB12 vector in the expression and purification of *Eg*PAAS, this enzyme was ultimately only partially purified with significant contamination of *E. coli* chaperone proteins.

### Protein Crystallization and Structural Determination

Crystals for the various plant AAADs were grown at 4 °C by hanging-drop vapor diffusion method with the drop containing 0.9 µL of protein sample and 0.9 µL of reservoir solution at a reservoir solution volume of 500 µL. The crystallization buffer for the *At*PAAS contained 0.16 M ammonium sulfate 0.8M HEPES:NaOH pH 7.5 and 20%w/v PEG 3350. Crystals were soaked in a well solution containing 15 mM L-phenylalanine for six hours and cryogenized with an additional 10% weight/volume ethylene glycol. *Ps*TyDC crystals were formed in 1.2 M ammonium Sulfate 0.1 Bis Tris pH 5.0 and 1% w/v PEG 3350. Crystals were soaked in the presence of 4mM L-tyrosine for 12 hours and cryoprotected with an additional 25% weight/volume ethylene glycol. 0.22 M calcium chloride and 12% w/v PEG 3350 formed the *Cr*TDC crystals which were subsequently soaked with 10mM L-tryptophan for 16 hours and then cryogenized with an additional 18% weight/volume ethylene glycol. Finally, to form the *Rr*4HPAAS crystals, protein solution was mixed with a reservoir buffer of 0.21 M potassium thiocyanate and 22% w/v PEG 3350. Ligand soaks for this crystal proved unsuccessful and ultimately the crystals were cryoprotected with an additional 13% weight/volume PEG 3350 in the absence of ligand. The *Ps*TyDC structure was determined first by molecular replacement using the insect DDC structure (58) as the search model in Molrep (59). The resulting model was iteratively refined using Refmac 5.2 (60) and then manually refined in Coot 0.7.1 (61). The *Cr*TDC, *At*PAAS and *Rr*4HPAAS structures were solved by molecular replacement using the refined *Ps*TyDC structure as the search model, followed by refinement procedure as described above.

### Enzyme assays

The *in vitro* decarboxylation and aldehyde synthase activities of the wild-type *Ps*TyDC, *Ps*TyDC^H205N^ and *Ps*TyDC^Y350F^ were assayed in 100 μL reaction buffer containing 50 mM Tris, pH 8.0, 100 μM PLP, 0.5 mM L-tyrosine, and 20 μg of recombinant enzyme. Reactions were incubated at 30 °C for various time points and subsequently stopped in the linear range of product formation with 200 μL methanol. After clarification, the soluble fraction was analyzed by LC-MS-UV. Chromatographic separation and measurement of absorption at 280 nm were performed by an Ultimate 3000 liquid chromatography system (Dionex), equipped with a 150 mm C18 Column (Kinetex 2.6 µm silica core shell C18 100 Å pore, Phenomenex) and coupled to an UltiMate 3000 diode-array detector (DAD) in-line UV-Vis spectrophotometer (Dionex). Compounds were separated through the use of an isocratic mobile phase as previously described (57). The reduction of aldehyde products was achieved by the addition of ethanol containing a saturating concentration of sodium borohydride. The *Eg*PAAS enzyme assays were started by adding 2 µg of recombinant protein into 200 µL reaction buffer containing 50 mM Tris pH 8.0, and 2 mM L-phenylalanine. Reactions were incubated for various time points at 30 °C, and the reactions were stopped with equal vol of 0.8 M formic acid, extracted with 150 µL of ethyl acetate and analyzed by gas chromatography–mass spectrometry (GC-MS) as previously described against an analytical phenylacetaldehyde standard (57). The initial substrate selectivity was measured through the detection of the hydrogen peroxide co-product using Pierce Quantitative Peroxide Assay Kit (Pierce) and a standard curve of hydrogen peroxide. Reactions were conducted as described using reaction mixtures containing 0.5 mM amino acid substrate concentrations. Triplicate reactions were stopped after five minutes of incubation at 30 °C with an equal volume of 0.8 M formic acid and measured by absorbance at 595 nm.

### Metabolic engineering and metabolic profiling of transgenic yeast by LC-HRAM-MS

*Li*AAS (Phytozome 12: *Lagerstroemia indica* scaffold RJNQ-2017655) and *Mm*AAS (Phytozome 12: *Medinilla magnifica* scaffold WWQZ-2007373) were synthesized as gBlocks (IDT) with *S. cerevisiae* codon optimization. Ectopic expression of various AAADs in *S. cerevisiae* was achieved through the use of p423TEF, a 2-μm plasmid with the *HIS3* auxotrophic growth marker for constitutive expression (62). 15 mL cultures of transgenic *S. cerevisiae* BY4743 strains were grown in 50 mL mini bioreactor tubes for 24 hours with shaking at 30 °C. The cultured cells were subsequently pelleted, washed, disrupted, and clarified for LC-HRAM-MS analysis as previously described (65). *PpDDC* (NP_744697.1), *PsNCS2* (AKH61498), and *BvTyH* (AJD87473) were synthesized as gBlocks (IDT) with *S. cerevisiae* codon optimization. PCR amplicons or gBlocks were ligated into the entry vector pYTK001 and subsequently assembled into 2-μm, *pTDH3*, *tTDH1*, *HIS3* multigene vectors for constitutive expression in *S. cerevisiae* (63). A second multigene vector, containing the *S. cerevisiae* tyrosine-metabolism feedback-resistant mutants *ARO4*^K229L^ and *ARO7*^G141S^, was additionally used to boost tyrosine flux as previously described (57). *S. cerevisiae* lines were transformed with various multi gene vectors to assay ectopic (S)-norcoclaurine production. Here, clarified media extracts from bioreactor cultures were diluted with an equal volume of 100% methanol and analyzed directly by LC-HRAM-MS. Raw MS data were processed and analyzed using MZmine2 (64). Data files were first filtered to only include positive mode ions above the noise filter of 1e5. Shoulder peaks were next removed using the FTMS shoulder peaks filter function. Chromatograms were assembled using the chromatogram builder function and smoothed using the peak smoothing function. Chromatograms were subsequently separated into individual peaks using chromatogram deconvolution. The resulting peak lists were aligned using the join aligner function and omitted peaks were identified and added using the gap filling function. The project parameters were then grouped by triplicate and the peak areas were exported for subsequent statistical analysis and graph generation.

### Molecular dynamics simulation and analysis

All simulations were performed using GROMACS 5.1.4 (65) and the CHARMM36 force field (66). The non-standard residue LLP was parameterized using Gaussian (67) and the Force Field Toolkit (FFTK) (68) implemented in VMD (69) based on the initial parameters provided by the CGenFF program (70–73). A number of *Cr*TDC residues buried deeply within the protein or at the monomer-monomer interface were modeled in their neutral forms based on PROPKA (74, 75) calculation results: Asp^268-A/B^, Asp^287-A/B^, Asp^397-A/B^, Lys^208-A/B^, and Glu^169-A^. All the histidines were kept neutral, with a proton placed on the ε-nitrogen except for His^203^ and His^318^, for which the proton was placed on the δ-nitrogen to optimize hydrogen bond network. All simulation systems were constructed as a dimer solvated in a dodecahedron water box with 0.1 M NaCl (Fig. S11) and a total number of atoms of ∼124,000. Prior to the production runs listed in Table S4, all systems were subjected to energy minimization followed by a 100-ps NVT and a 100-ps NPT runs with the protein heavy atoms constrained. In all simulations the van der Waals interactions were smoothly switched off from 10 Å to 12 Å. The electrostatic interactions were computed with the Particle-Mesh-Ewald (PME) method (76) with a grid spacing of 1.5 Å, a PME order of 6, and a cutoff of 12 Å in real space. The system temperature was kept at 300 K using the velocity-rescaling thermostat (77), and the system pressure was kept at 1 bar with the Parrinello-Rahman barostat (78, 79). All bonds were constrained using LINCS (80, 81) to allow an integration timestep of 2 fs. The helix-unfolding simulation was performed using the metadynamics method (82) as implemented in PLUMED (83). A 10-ns metadynamics simulation was performed on System 1 by placing gaussian potentials (height=35 kJ/mol, sigma=0.35 rad) every 500 steps on the collective variables, which were chosen as the backbone dihedral angles Ψ and Φ of residues 346 to 350. We should emphasize that this simulation was not intended for an accurate free energy calculation and instead was only used to generate an unfolded structure of the short helix (residues 346-350). The resulting unfolded large loop structure was then used in all systems, each of which was subjected to 12 replicas of 50-ns MD simulations listed in Table S4. Clustering analysis was performed using gmx cluster over all simulated trajectories with a RMSD cutoff of 3.5 Å. 3D occupancy maps were created at a resolution of 1 Å^3^ using the VMD VOLMAP plugin. DSSP calculations (84) were performed with gmx do_dssp implemented in GROMACS. An output of H (ɑ-helix), I (*π*-helix) or G (3_10_-helix) was considered as a helix and the corresponding residue was assigned a helical content of 1; otherwise a helical content of 0 was assigned. Clustering and occupancy analysis as well as the average helical content calculations were performed on the combined trajectories of all simulation replicas for a given *Cr*TDC system. The two monomers of a *Cr*TDC dimer were treated equivalently in these analyses. All simulation figures were made using VMD.

## Accession codes

The sequences of *P. somniferum*, *C. roseus*, *E. grandis* genes reported in this article are deposited into NCBI GenBank under the following accession numbers: *PsTyDC* (MG748690), *CrTDC* (MG748691), *EgPAAS* (MG786260). The atomic coordinates and structure factors for *Ps*TyDC, *Cr*TDC*, Rr*4HPAAS, and *At*PAAS have been deposited in the Protein Data Bank under the accession numbers 6EEM, 6EEW, 6EEQ and 6EEI, respectively.

## Acknowledgments

This work was supported by the Pew Scholar Program in the Biomedical Sciences (J.K.W.), the Searle Scholars Program (J.K.W.), the National Science Foundation (CHE-1709616, J.K.W.), the Keck Foundation (J.K.W.), and direct grants from the Chinese University of Hong Kong (Y.W.). This work is based on research conducted at the Northeastern Collaborative Access Team (NE-CAT) beamlines, which are funded by the National Institute of General Medical Sciences from the National Institutes of Health (P41 GM103403). The Pilatus 6M detector on NE-CAT 24-ID-C beam line is funded by a NIH-ORIP HEI grant (S10 RR029205). This research used resources of the Advanced Photon Source, a U.S. Department of Energy (DOE) Office of Science User Facility operated for the DOE Office of Science by Argonne National Laboratory under Contract No. DE-AC02-06CH11357.

## Author Contributions

M.P.T.S. and J.K.W. conceived the idea and designed the research. M.P.T.S., T.S, and M.A.V. performed cloning and phylogenetic analysis. M.P.T.S. conducted enzyme purification, enzyme assays, crystallization, and structural elucidation. Y.C and Y. W performed the molecular dynamic simulations and data interpretation. M.P.T.S., Y.C, Y.W. and J.K.W. wrote the paper.

## Supplementary Information

### Supplementary Figures

**Fig. S1.**
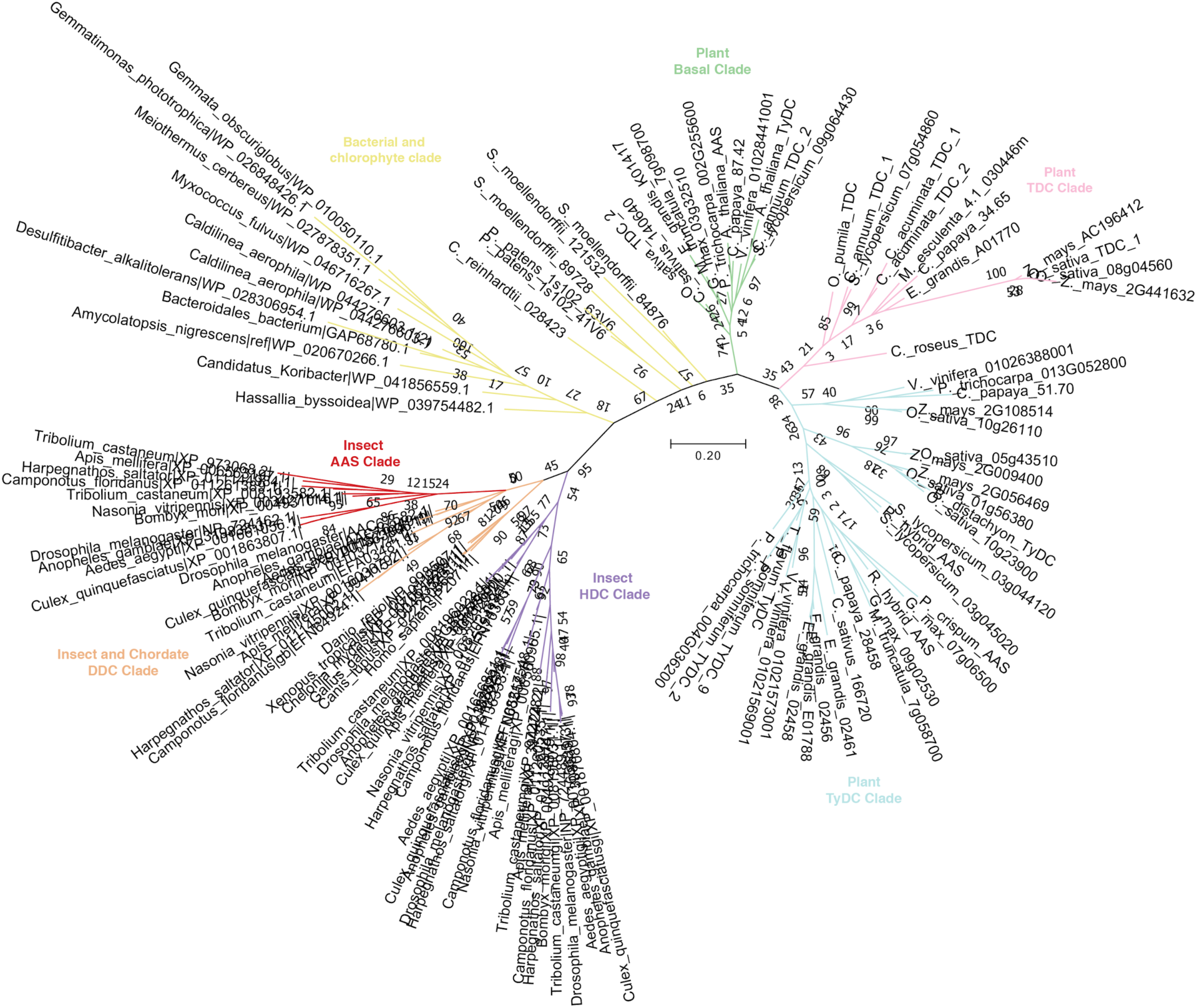
A maximum-likelihood phylogenetic tree of select chordata, insect, bacterial, and plant AAADs. This tree is populated with sequences from all Phytozome V12 species, all attainable characterized NCBI plant AAAD sequences and select eubacteria, chordata, and insect NCBI sequences. Bootstrap values are indicated at the tree nodes. The bootstrap consensus unrooted trees were inferred from 500 replicates. The scale measures evolutionary distances in substitutions per amino acid. The green, pink and blue branches correspond to the basal, TDC and TyDC plant clades, respectively. The yellow branches correspond to the chlorophytes and bacterial AAAD sequences. The purple clade corresponds to insect histidine decarboxylase (HDC) sequences, while the orange clade represents insect and chordata DDC sequences. The red insect AAS clade emerged from the insect DDC clade and contains asparagine substitution in place of the typically considered active-site histidine. The evolutionary history of these enzymes indicates that animal and plant AAADs are monophyletic and have evolved independently from a universal common ancestor.

**Fig. S2.**
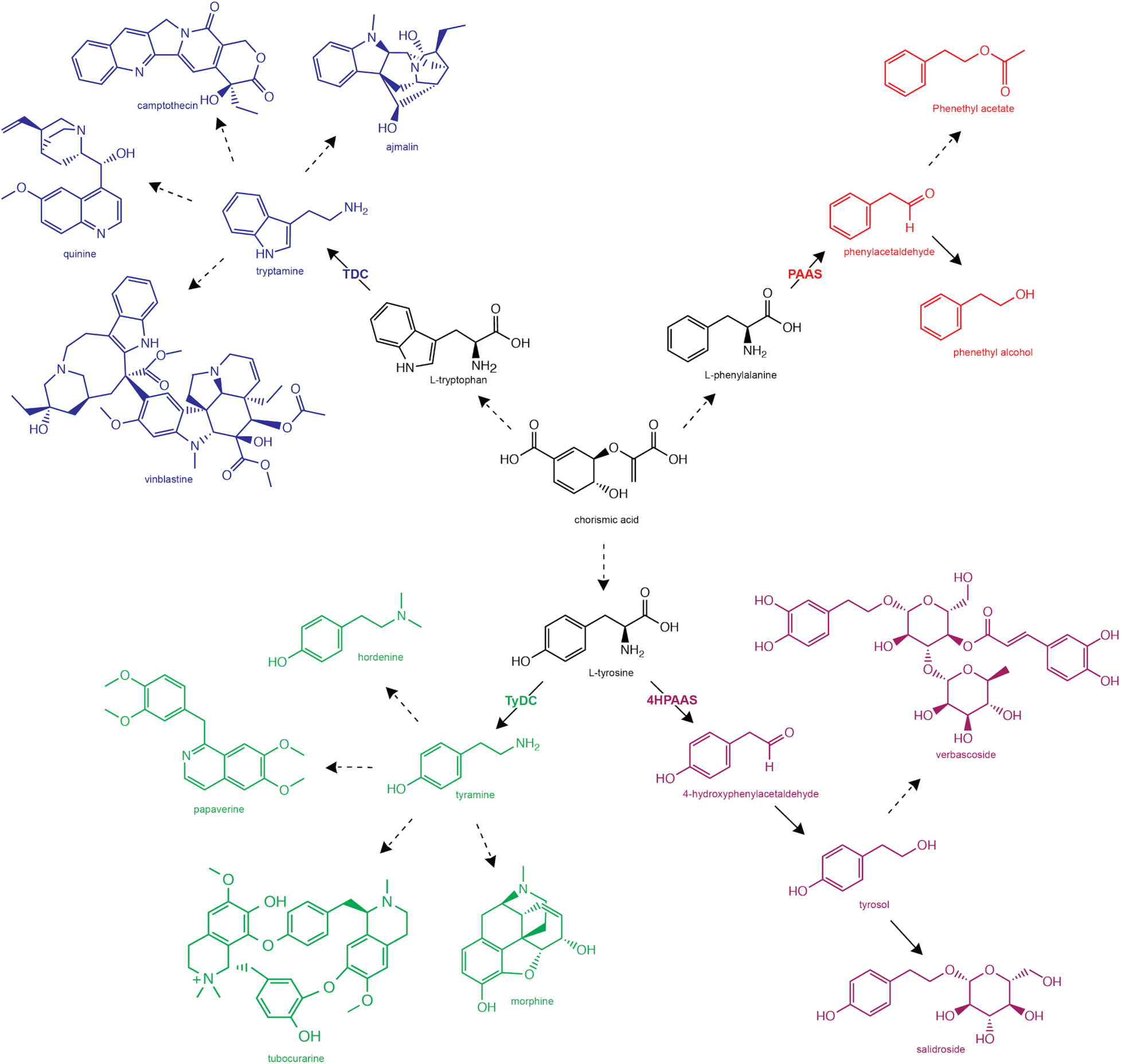
Diverse specialized metabolic pathways downstream of various plant AAADs. In plants, the three proteinegic L-aromatic amino acids, L-tryptophan, L-tyrosine, and L-phenylalanine are all downstream of chorismate derived from the shikimate pathway. From these primary metabolites, plant AAADs catalyze the first biotransformation in divergent specialized metabolic pathways. The TDC enzyme and some downstream products are shown in blue, the PAAS and several downstream products are shown in red, the TyDC and select downstream products are shown in green, and the 4HPAAS and a few downstream products are shown in purple. Solid arrows represent single enzyme catalyzed reactions while the dotted arrows indicate multiple enzymatic steps.

**Fig. S3.**
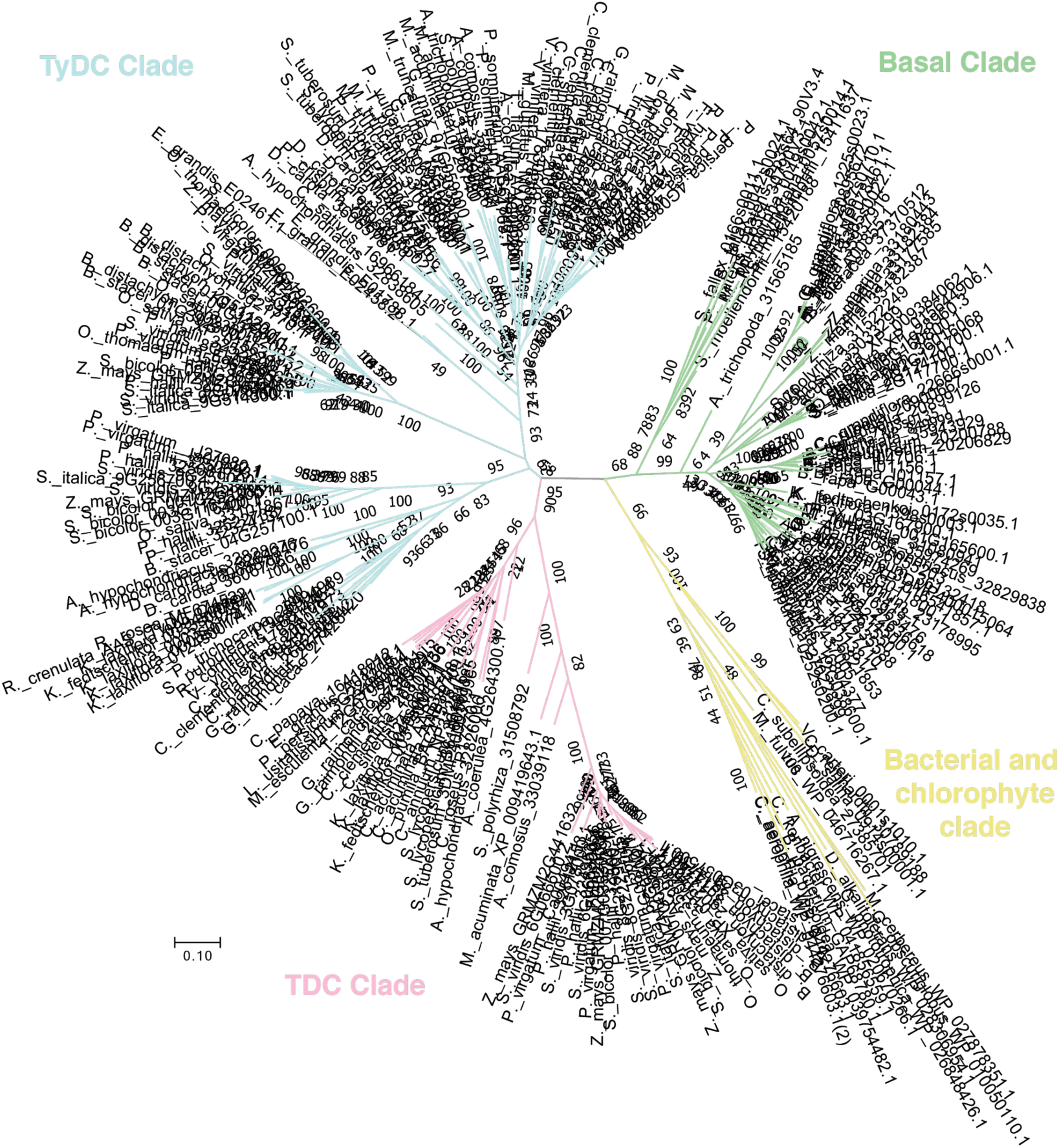
A maximum-likelihood phylogenetic tree of AAADs. This tree is populated with sequences from all Phytozome V12 species, all attainable characterized plant AAAD sequences and select eubacteria from NCBI. Bootstrap values are indicated at the tree nodes. The bootstrap consensus unrooted trees were inferred from 500 replicates. The scale measures evolutionary distances in substitutions per amino acid. Green, pink and blue branches correspond to the basal, TDC and TyDC clades, respectively. The yellow branches correspond to the chlorophytes and bacterial AAAD sequences. The plant AAAD clades were annotated according to their relation to ancestral sequences (the basal clade is most closely related to bacterial and chlorophyte AAADs) and their apparent substrate selectivity (the TDC clade contains a number of characterized enzymes with exclusive indolic substrate specificity, while the TyDC clade is represented by characterized enzymes with phenolic substrate selectivity). While the basal and TyDC clades contain substitutions impliciative of AAS chemistry, these mutations occur independently and sporadically through plant taxonomy.

**Fig. S4.**
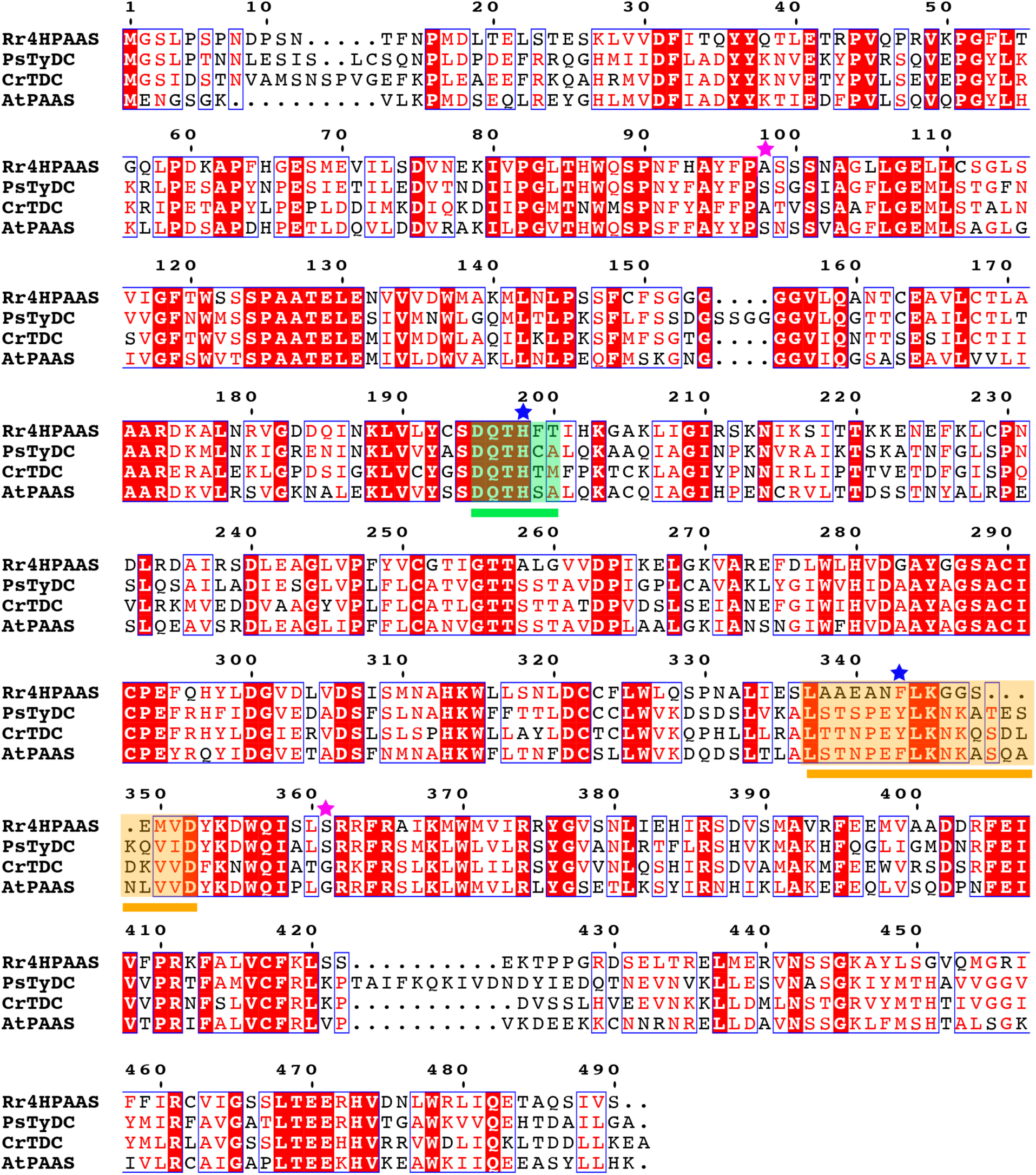
Multiple sequence alignment of four crystallized plant AAADs. The short catalytic loop is highlighted and underlined in green and the large catalytic loop is highlighted and underlined in orange. The substrate selectivity residues are marked with pink stars and the catalytic mechanism dictating residues are marked with blue stars. The multiple sequence alignment was generated with ClustalW2 (1) and displayed with ESPript 3.0 (2).

**Fig. S5.**
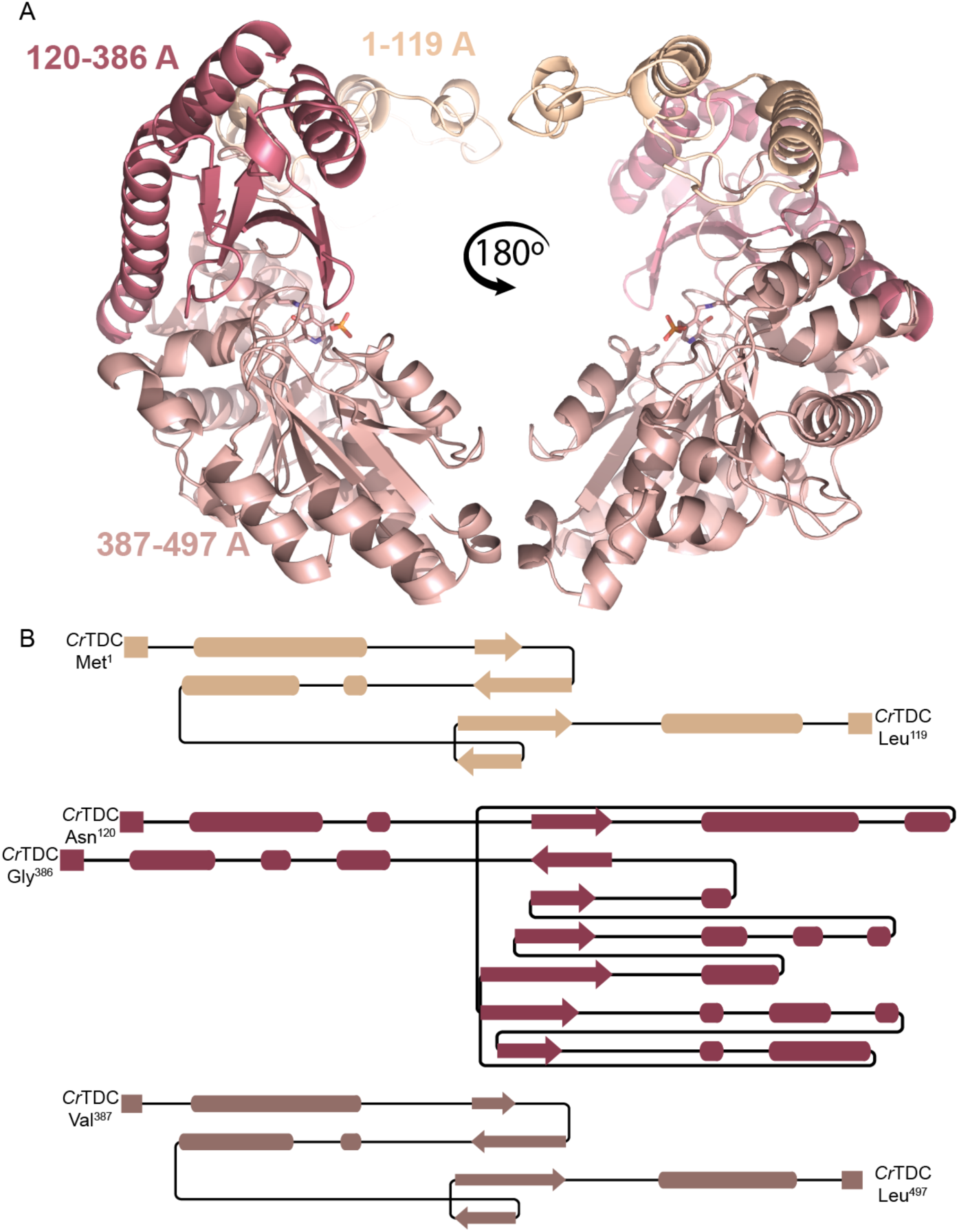
Topology of plant AAADs as observed in *Cr*TDC. (*A*) Plant AAAD segments as displayed by the *Cr*TDC structure. Each monomer is composed of the N-terminal *Cr*TDC^1-119^ (beige), middle *Cr*TDC^120-386^ (maroon) and C-terminal *C*rTDC^387-497^ (salmon) segments. (*B*) Topology diagram for each of the three *Cr*TDC segments. The segment diagrams were generated though Pro-origami using DSSP secondary structure program (3).

**Fig. S6.**
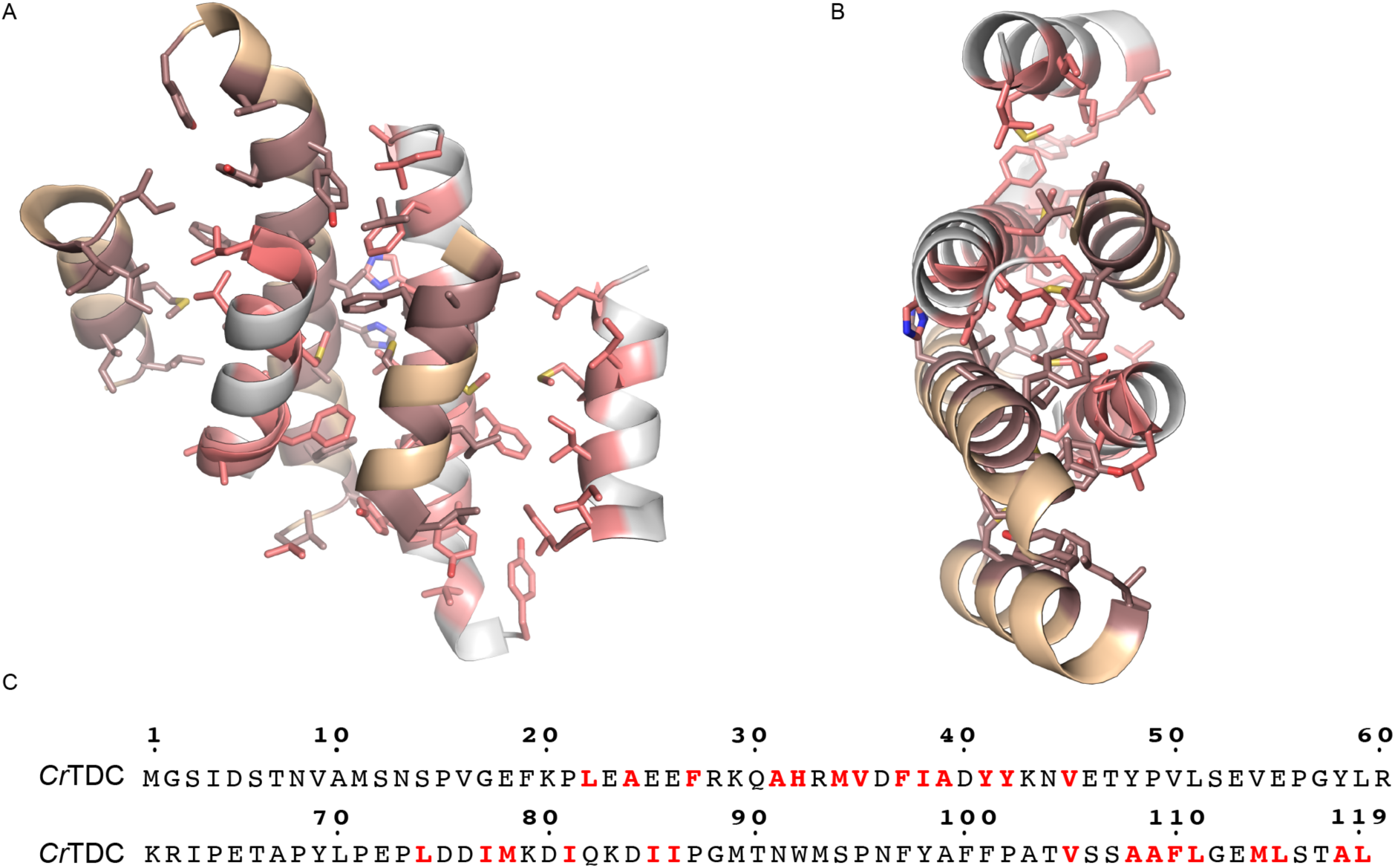
Intermonomer association of N-terminal segments from two *Cr*TDC chains. Side (*A*) and top (*B*) views of the aromatic and hydrophobic residues forming the intermolecular interaction of the *Cr*TDC homodimer. One monomer is colored in orange with maroon hydrophobic or aromatic residues, whereas the second monomer is colored in white with pink hydrophobic or aromatic residues. (*C*) Sequence of the *Cr*TDC N-terminal segment with hydrophobic or aromatic residues involved in intermonomer association highlighted in red.

**Fig. S7.**
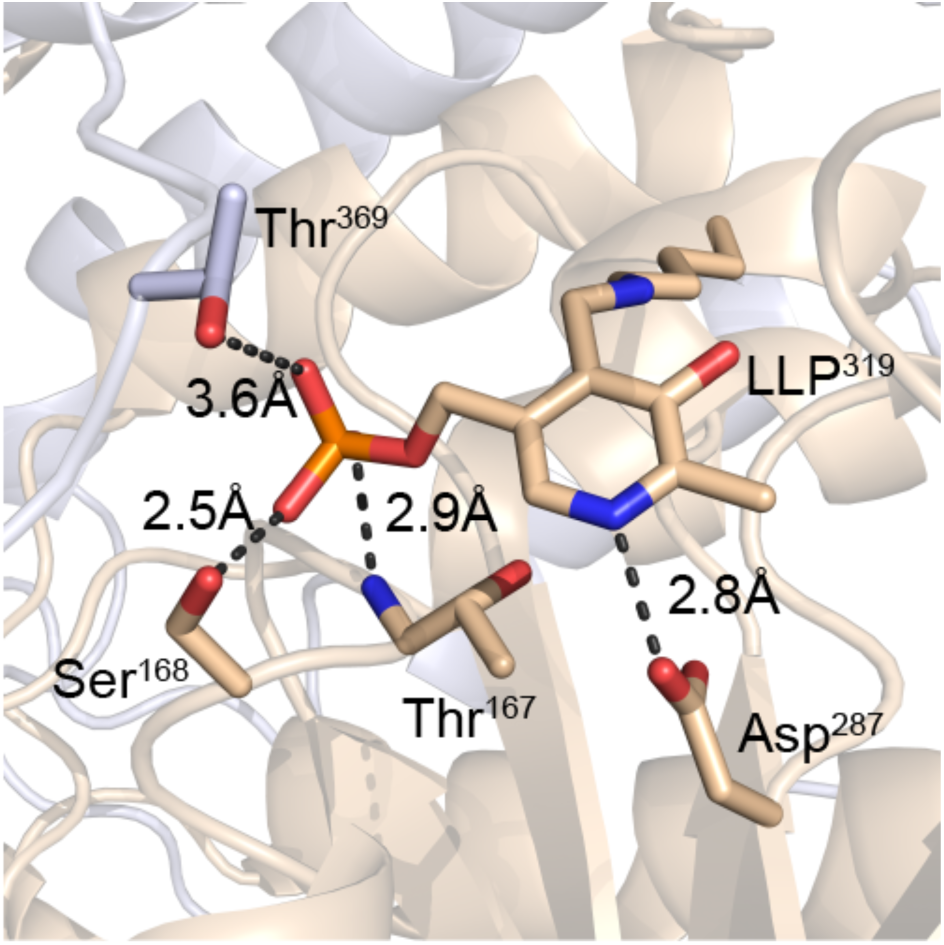
Coordination of LLP by the *Cr*TDC active-site residues. Cartoon and stick representation of LLP coordination in *Cr*TDC. Chain A is colored in beige and chain B is colored in white.

**Fig. S8.**
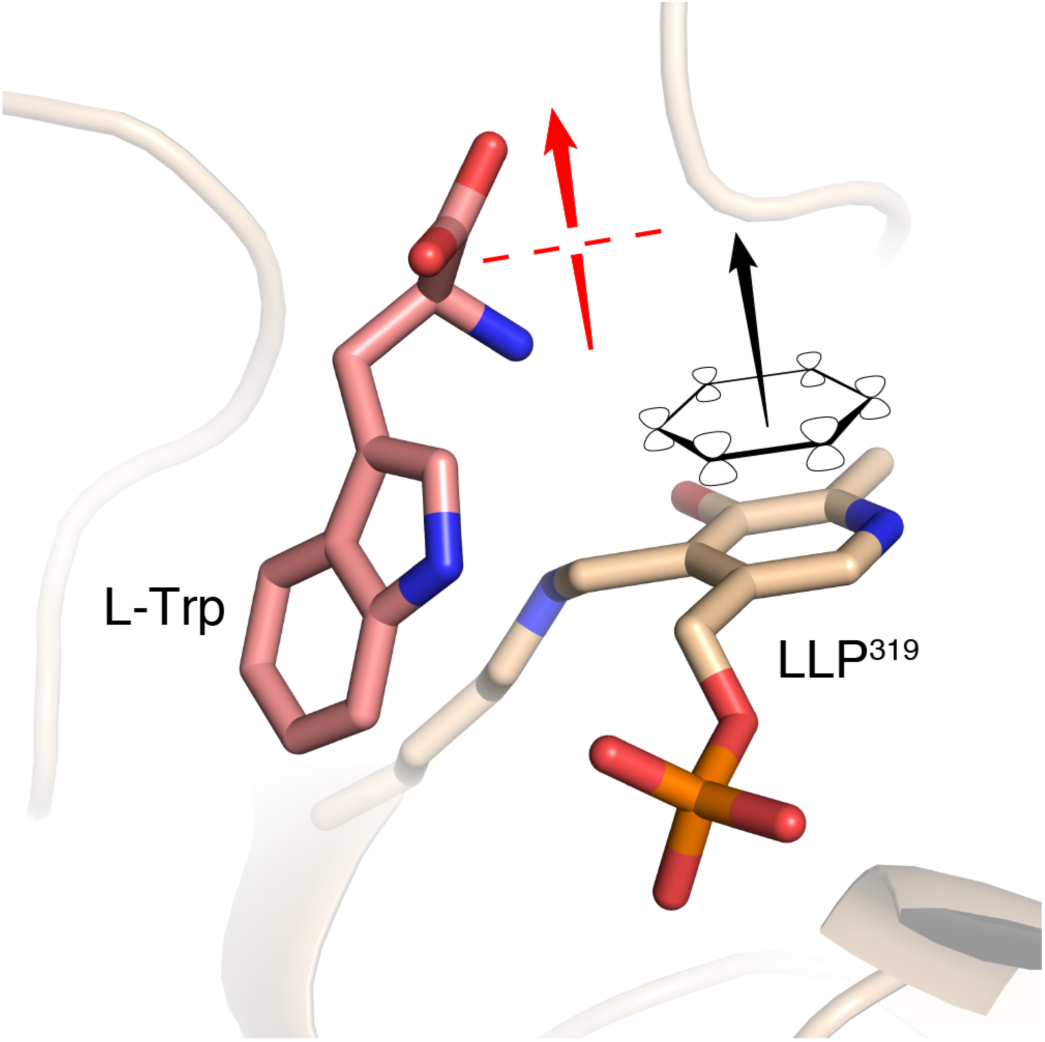
Orientation of the alpha carbon carbonyl bond relative to the plane of the pyridoxal imine system. As per the Dunathan hypothesis, PLP enzymes exhibit stereospecific cleavage of bonds orthogonal to the pyridine ring pi-system electrons (shown as black ring and arrow) (4). In the case of PLP decarboxylases, the alpha carbon carbonyl bond of the substrate is positioned perpendicular to the plane of the pyridine ring (shown as red arrow).

**Fig. S9.**
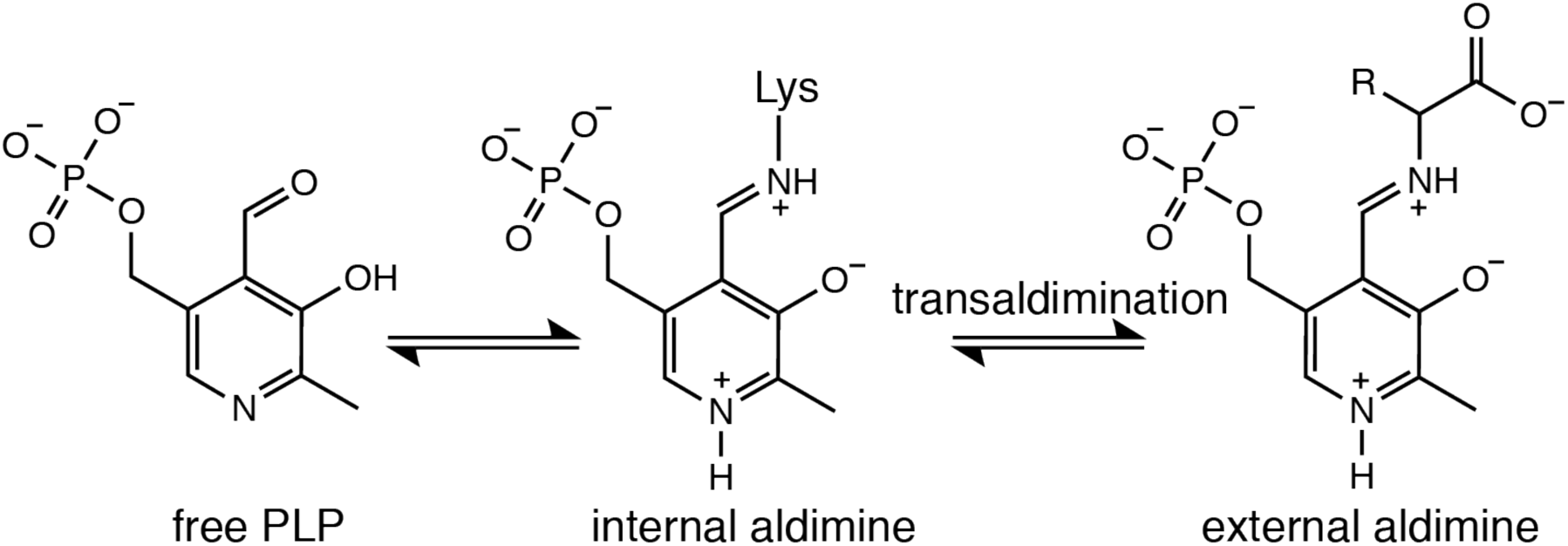
Schematic of the formation of PLP internal and external aldamines. First, the internal aldimine is formed when the aldehyde group of the PLP coenzyme forms an imine with the conserved active site lysine. Second, the external aldimine is formed upon the imine exchange between the ζ-amino group of the lysine and the α-carbon amine of the substrate.

**Fig. S10.**
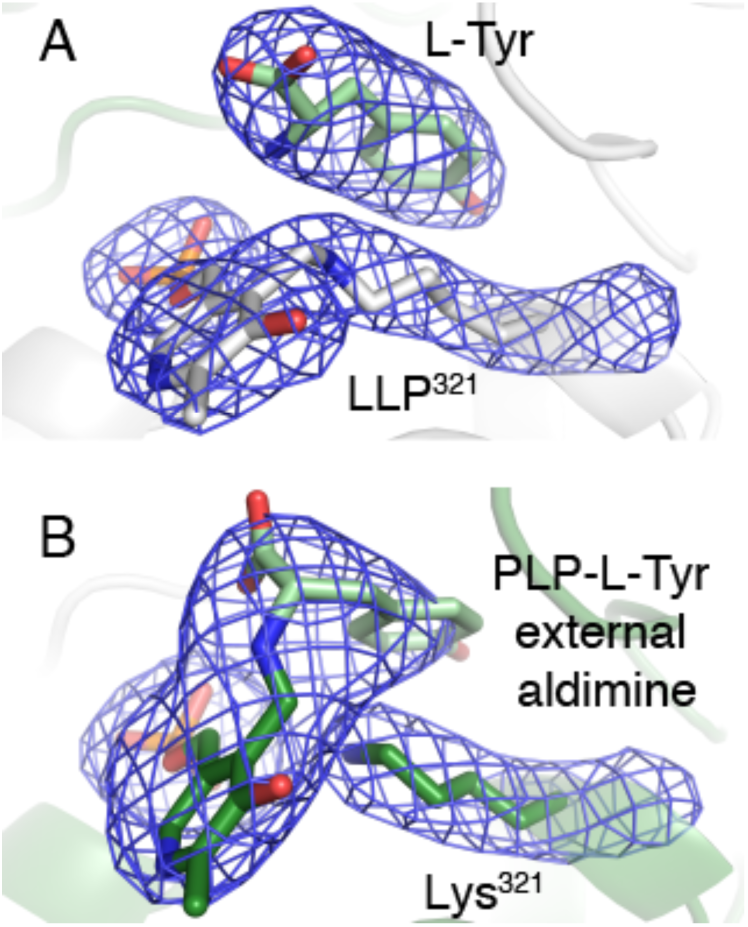
Transaldimination as captured by two active sites of the *Ps*TyDC homodimer. (*A*) The PLP-Lys^321^ internal aldimine as modeled in one of the active sites of the *Ps*TyDC homodimer. (*B*) The PLP-L-tyrosine external aldimine as captured by the other active site of the *Ps*TyDC homodimer. The Chain A and Chain B are colored in green in gray, respectively, and the |2*Fo – Fc*| electron density map is contoured at 2 σ.

**Fig. S11.**
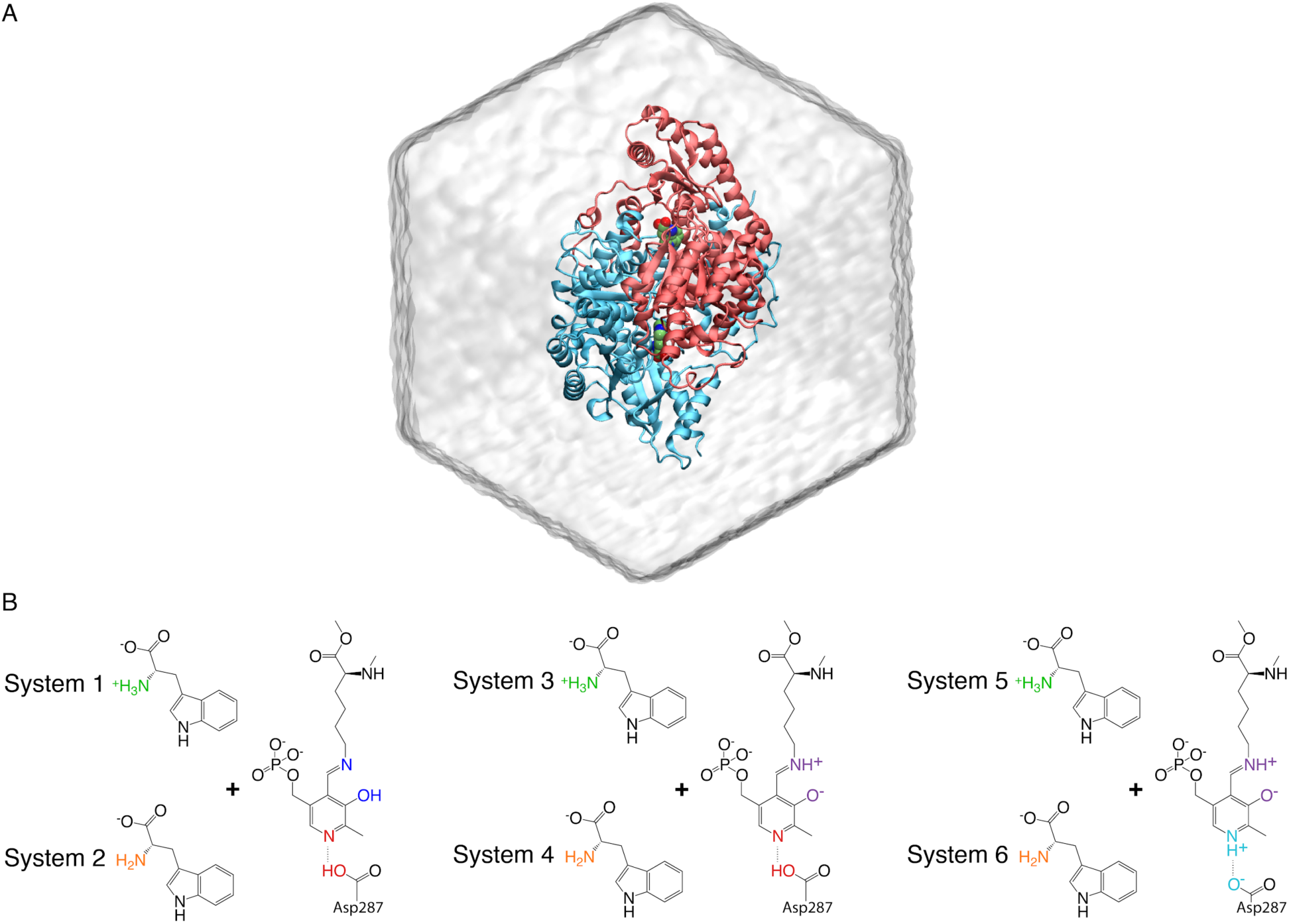
MD simulation systems of holo-*Cr*TDC with LLP and L-tryptophan in different protonation states. (*A*) The dodecahedron simulation box with the two monomers of *Cr*TDC colored in red and blue, respectively. Water molecules are shown as transparent surfaces. (*B*) Six holo-*Cr*TDC systems simulated in this work.

**Fig. S12.**
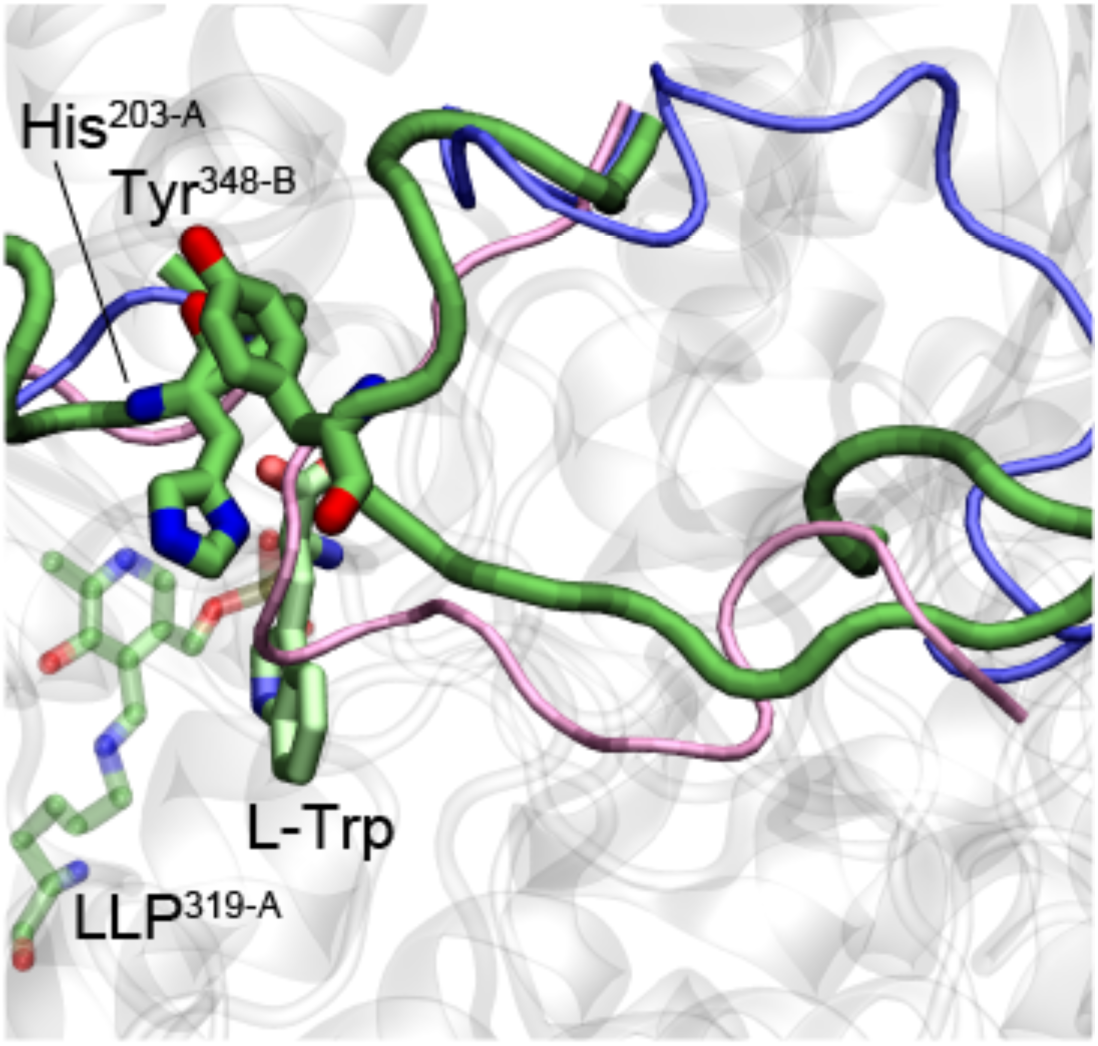
*Cr*TDC MD semi-closed conformation. A snapshot from the MD simulation of *Cr*TDC System 1 at t=398 ns, exhibiting a semi-closed loop conformation. The open and closed conformations of the loops observed from the crystal structures are shown in blue and pink tubes, respectively.

**Fig. S13.**
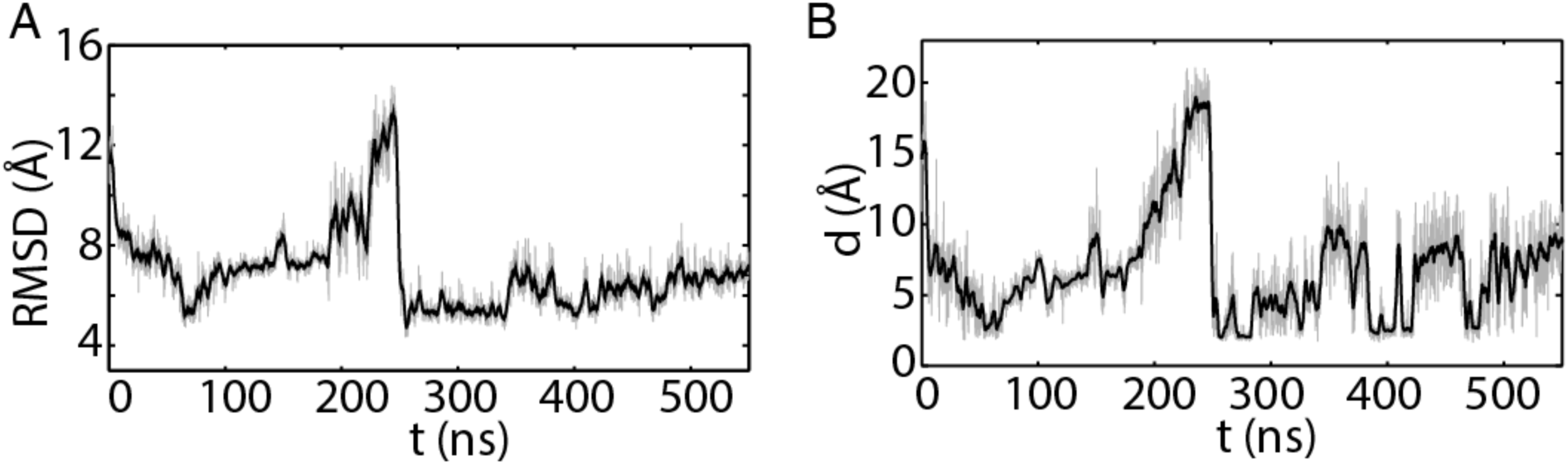
Large loop conformation as measured by RMSD and atomistic distances in the 550-ns simulation of *Cr*TDC system 1. (*A*) RMSD of large loop C_α_ atoms with respect to the modeled closed-state *Cr*TDC. (*B*) The minimal distance between Tyr^348-B^ and His^203-A^. Black curves represent running averages (window size: 101) performed on data colored in gray.

**Fig. S14.**
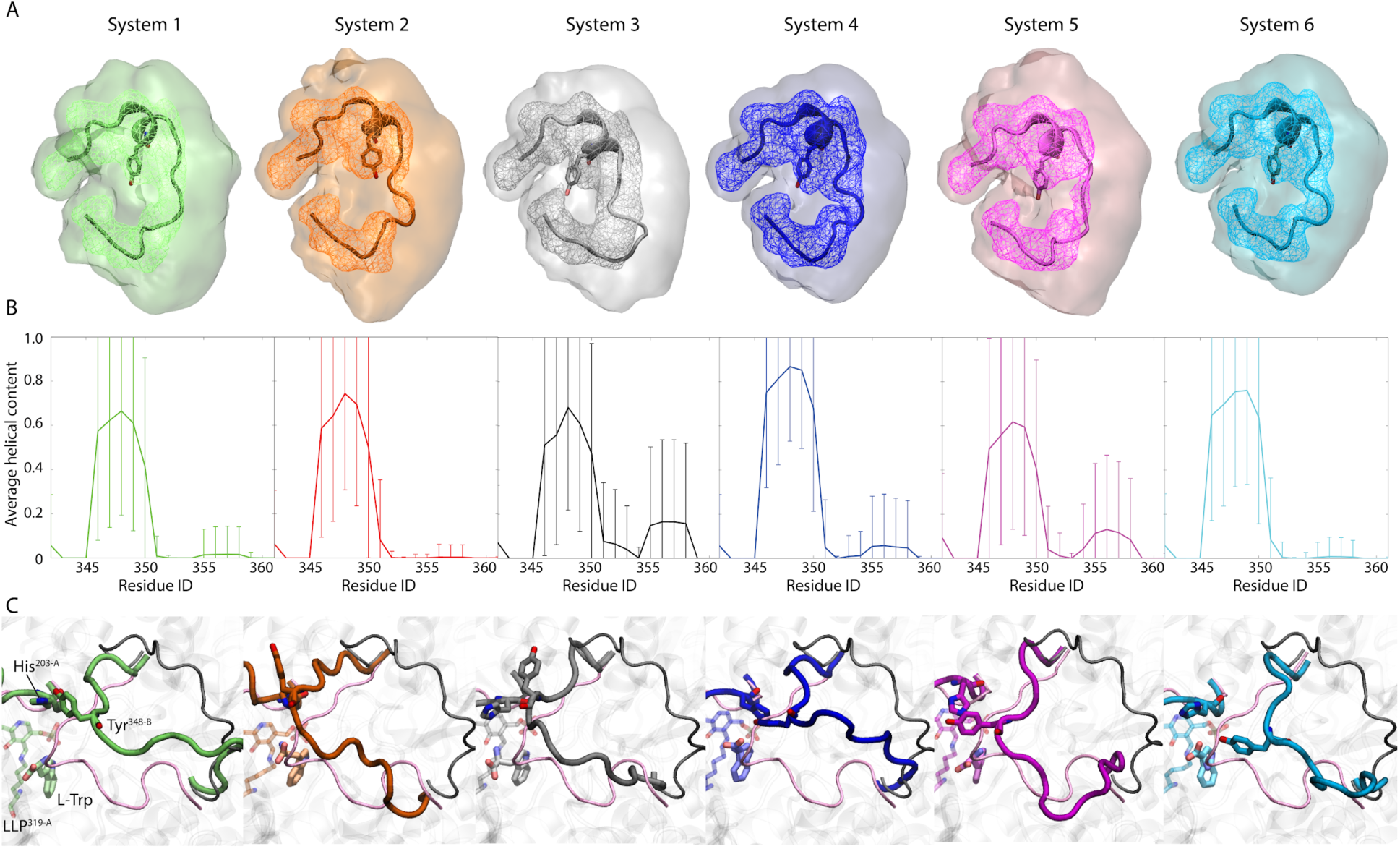
Large loop conformations revealed by MD simulations of holo-*Cr*TDC. (*A*) Clustering analysis and occupancy calculation results performed on the six replicas of 100-ns simulations of each *Cr*TDC system. Centroid structure of the largest cluster from clustering analysis is shown in Cartoon representation, where a short helix (residues 346-350) is seen across all systems. Isosurfaces of 30% and 1% occupancy are shown in wireframes and transparent surfaces, respectively. (*B*) Average helical content of the large loop in the simulations described in (*A*). Error bars indicate standard deviations. (*C*) Snapshots from selected 50-ns simulations of *Cr*TDC systems 1-6 initiated with the short helix in an unfolded state (Table S4). Initial structure of the large loop in this unfolded state is shown in black thin tube, with the closed conformations of the loops from crystal structure shown in pink thin tubes.

**Fig. S15.**
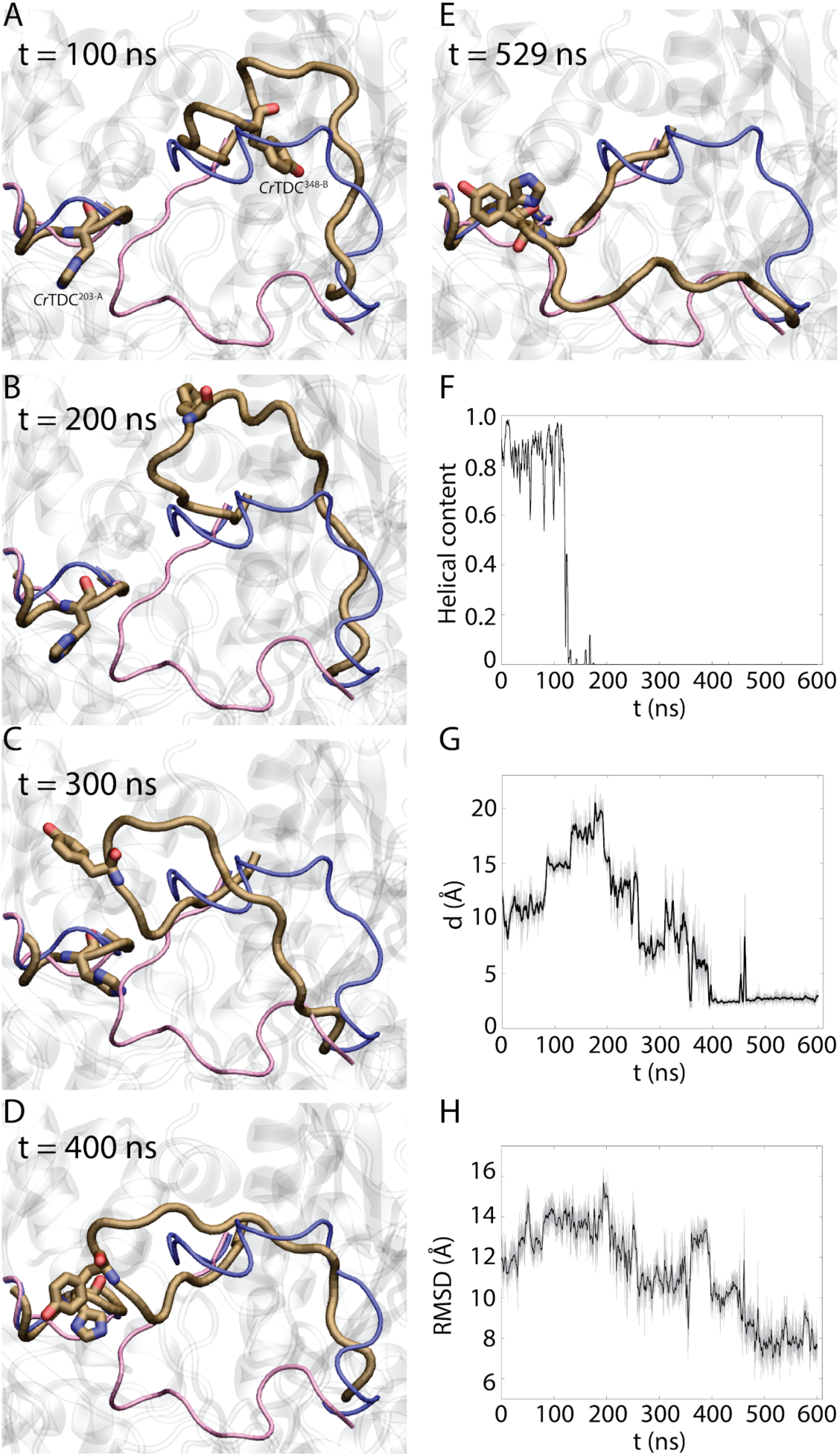
Large loop conformations revealed by a 600-ns MD simulation of apo-*Cr*TDC. (*A-E*) Simulation snapshots with residues Tyr^348-B^ and His^203-A^ highlighted. Loop conformations in the open and the modeled closed-state *Cr*TDC are colored in blue and red, respectively. (*F*) Helical content of the large loop during the 600-ns apo-simulation. Note that the loss of helical content precedes the large-scale loop closing motion shown in (*A-E*). (*G*) Minimal distance between His^203-A^ and Tyr^348-B^. (*H*) C_ɑ_ RMSD of the large loop with respect to the modeled closed-state *Cr*TDC.

**Fig. S16.**
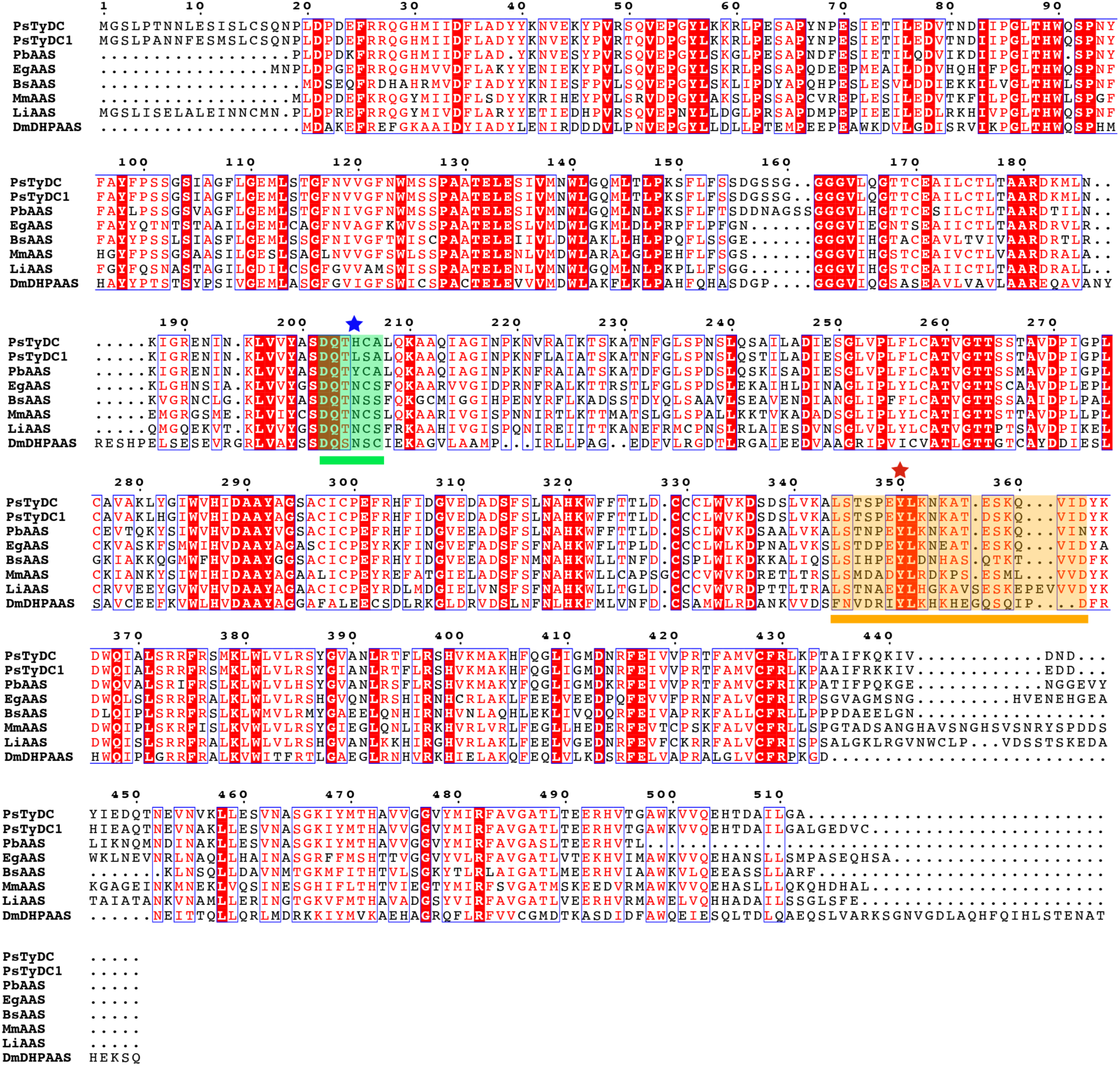
Multiple sequence alignment of plant and insect AASs together with *Ps*TyDC highlighting naturally occurring substitutions at the small-loop histidine. The short catalytic loop is highlighted and underlined in green. The small-loop histidine conserved among canonical decarboxylases is labeled with a blue star. Variation in this residue may be implicative of alternative reaction mechanisms. The large loop is highlighted and underlined in orange. Although all the sequences display the red-stared catalytic tyrosine typically conserved in decarboxylation chemistry, *Eg*PAAS and *Drosophila melanogaster* 3,4-dihydroxyphenylacetaldehyde synthase (*Dm*DHPAAS) (5) display confirmed aldehyde synthase activity. *Ps*TyDC is crystallized in this study, while *Ps*TyDC1 (NCBI accession P54768) displays a small-loop histidine substitution but was previously annotated as a decarboxylase (6). *Papaver bracteatum* scaffold TMWO-2021544 (*Pb*AAS), B*egonia sp*. scaffold OCTM-2012326 (*Bs*AAS), *Medinilla magnifica* scaffold WWQZ-2007373 (*Mm*AAS), and *Lagerstroemia indica* scaffold RJNQ-2017655 (*Li*AAS) all contain small-loop His-to-Asn substitution characteristic of insect AASs. The multiple sequence alignment was generated using ClustalW2 (1) and displayed with ESPript 3.0 (2).

**Fig. S17.**
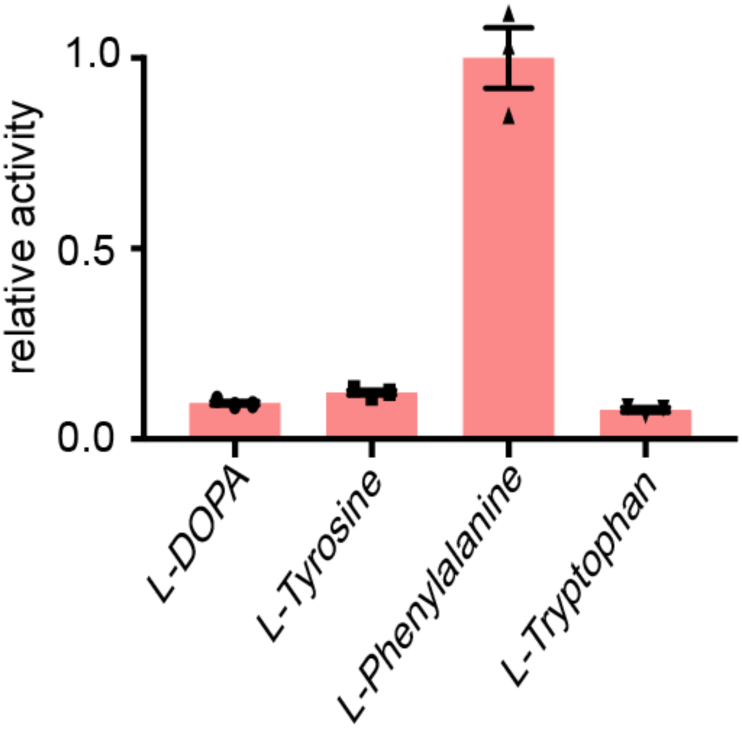
Relative selectivity of *Eg*PAAS towards various aromatic L-amino acid substrates. AAS activity of *Eg*PAAS was measured against various L-aromatic amino acid substrates at 0.5 mM substrate concentration. The relative activity was quantified through detection of the hydrogen peroxide co-product using the Pierce Quantitative Peroxide Assay Kit (Pierce). Error bars indicate standard error of the mean (SEM) based on biological triplicates.

**Fig. S18.**
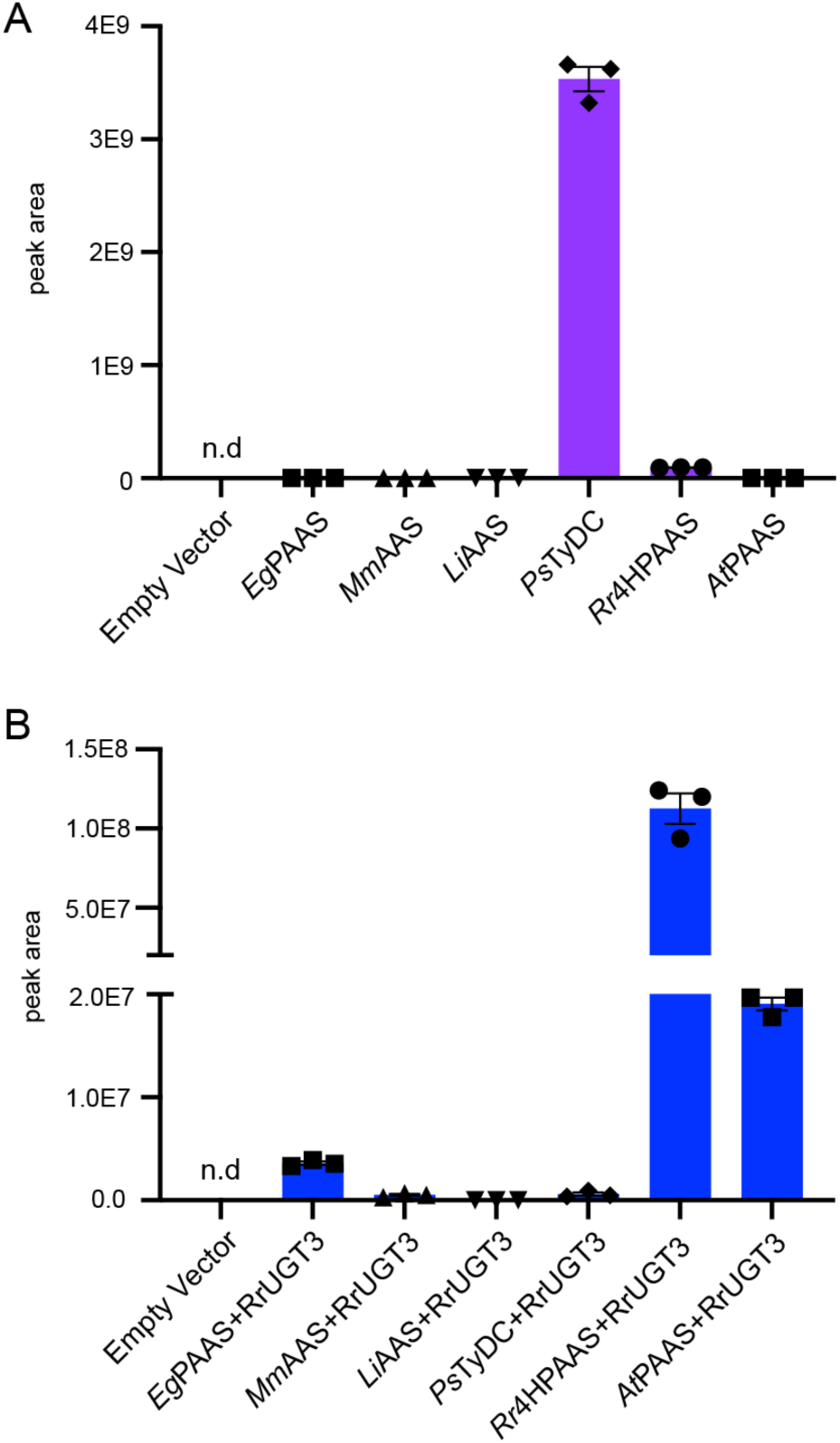
Relative canonical AAAD activity and AAS activity of various plant AAADs as assessed by tyramine and icariside D2 production, respectively, in transgenic yeast. (*A*) Tyramine accumulation from *in vivo* decarboxylation of L-tyrosine by various plant AAAD proteins in transgenic yeast. *Eg*PAAS, *Mm*AAS (Phytozome: *M. magnifica* scaffold-WWQZ-2007373), and *Li*AAS (Phytozome: *L. indica* scaffold-RJNQ-2017655) contain the signature His-to-Asn substitution as observed in insect AASs. (*B*) Icariside D2 accumulation in yeast expressing various plant AAADs alongside the *R. rosea Rr*UGT3 required for the 4-*O*-glycosylation of tyrosol (7). The 4-hydroxyphenylacetaldehyde product of 4HPAAS is reduced endogenously in yeast to form tyrosol prior to 4-*O*-glycosylation (7). Note that phenylacetaldehyde and phenylethyl alcohol are highly volatile, therefore could not be retained in yeast culture to be measured by LC-MS. Therefore, the relative canonical AAAD activity and AAS activity of these enzymes were assessed against the substrate L-tyrosine. The addition of *Rr*UGT3 facilitates the more reliable LC-MS detection of the AAS activity. Error bars indicate standard error of the mean (SEM) based on biological triplicates.

**Fig. S19.**
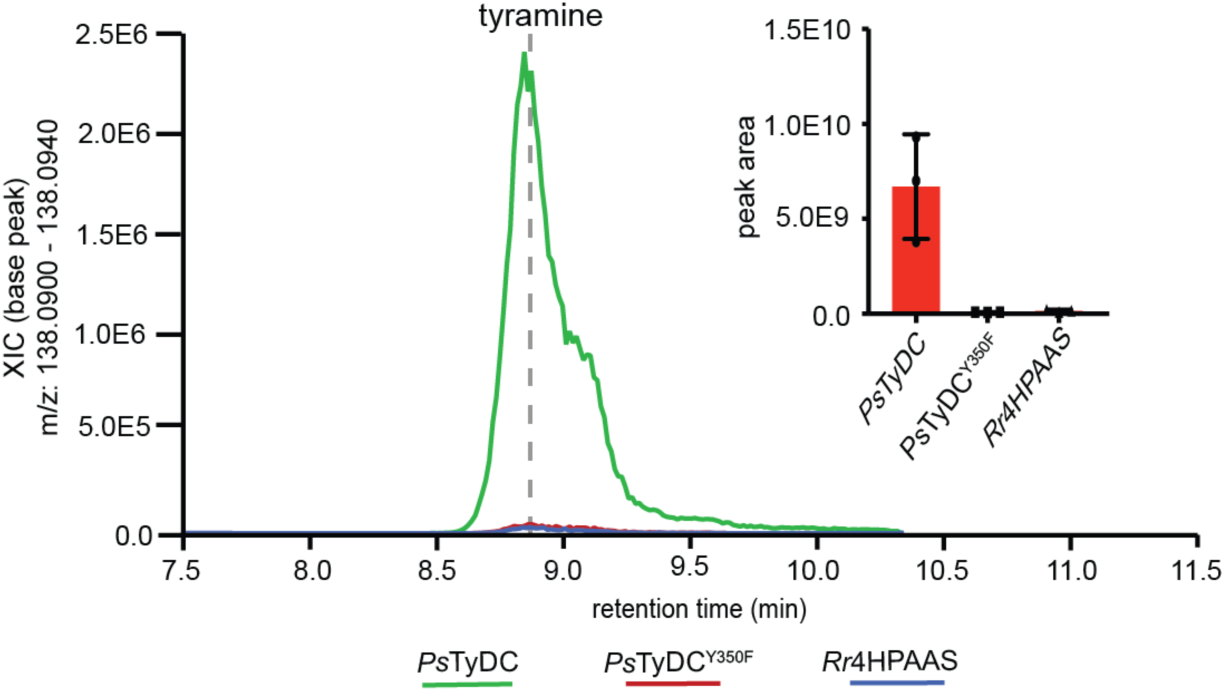
Relative *in vivo* L-tyrosine decarboxylase activity of various AAAD proteins in transgenic yeast. Yeast transformed with the multi-gene vector containing the wild-type *PsTyDC* produces tyramine rather than (S)-norcoclaurine. The multi-gene vectors used to transform yeast contain the requisite *PpDDC*, *PsNCS* and *BvTyH* in addition to either *PsTyDC*, *PsTyDC*^Y350F^ or *Rr4HPAAS*. Error bars indicate standard error of the mean (SEM) based on biological triplicates.

**Supplementary Video. Trajectory of a 550-ns simulation of *Cr*TDC System 1.** The large loop reached a semi-closed state during this simulation, with Tyr^348-B^ and His^203-A^ in frequent contact. For visualization clarity, water molecules and a large part of *Cr*TDC were not shown. Image smoothing was performed with a window size of 5 frames, which may have produced slight distortion of certain structures.

### Supplementary Notes

#### Supplementary Note 1

The residue range defined for the catalytic loops covers the area lacking significant secondary structure in the final *Cr*TDC model. The homologous sequence for the large loop in *Cr*TDC, *Ps*TyDC, *At*PAAS and *Rr*4HPAAS corresponds to residues 342-361, 344-363, 332-351 and 337-352, respectively. The small loop is represented in the *Cr*TDC, *Ps*TyDC, *At*PAAS and *Rr*4HPAAS sequences by residues 200-205, 202-207,190-195 and 195-200, respectively. In the CrTDC structure, the open conformation large loop lies on top of the upstream anchoring helix (*Cr*TDC residue range 335-341). This is particularly notable as the open conformation of this loop has not been observed in previously solved homologs. The large loop structure includes a single two turn helix containing the catalytic tyrosine. This loop helix interacts minimally with the preceding helix displaying tentative ionic interactions with Arg^340^. A similar two turn helix is not observed in the large loop of the closed conformation *Ps*TyDC structure, possibly due to the missing electron density of *Ps*TyDC residues 354-359. The paralogous human HDC, however, displays a homologous two turn helix in the closed conformation suggest that the secondary structure of this loop may be important throughout the conformational change. The B-chain of the *At*PAAS structure displays a partially modeled catalytic loop in the open conformation. Residues 339-445 were not built into this catalytic loop as this sequence range displayed poor electron density. Likewise, the majority of the large loop was not modeled in the *Rr*4HPAAS structure as there was poor electron density support.

#### Supplementary Note 2

The lysine-conjugated coenzyme PLP is simulated in either the enolimine (systems 1-2) or the ketoenamine form (systems 3-6) shown in Fig. S11. In aqueous solution, PLP aldimine is known to undergo reversible proton tautomerism between these two forms (8–10). Although when conjugated with an enzyme, the ketoenamine form is believed to dominate (8, 9), we decided to simulate both forms for the sake of completeness. The electron density map of *Cr*TDC suggests an electron shared between the pyridine nitrogen of LLP and Asp^287^. While PROPKA calculation supports a protonated Asp^287^, a deprotonated Asp^287^ is known to stabilize LLP with its pyridine nitrogen protonated (11–13). Therefore, while we modeled the enolimine form of LLP with Asp^287^ protonated (systems 1-2), both states of this residue were modeled in the ketoenamine form of the coenzyme (systems 3-6). The substrate L-tryptophan, which is a zwitterion at pH=7, is expected to lose the proton on its amine group prior to the formation of the external aldimine. Given that it is unclear when such deprotonation process occurs, we simulated L-tryptophan in both forms (Fig. S11). Taken together, six holo-*Cr*TDC systems were constructed (Table S4) and six replicas of 100-ns simulations were initially launched for each system. Analysis of these simulation trajectories reveals highly similar dominating conformation of the large loop, represented by the centroid structure from the largest cluster shown in Fig. S14*A*. The cartesian space visited by loop residues as enclosed by the occupancy isosurfaces (Fig. S14*A*) as well as the average helical content of the large loop measured by the program DSSP (Fig. S14*B*) are also similar across all six systems. These results suggest that in its initial transition from an open to a semi-closed state, conformational change of the large loop is not dictated by LLP and L-tryptophan protonation states. Indeed, in one of the three replicas of 600-ns apo *Cr*TDC simulations where neither PLP nor L-tryptophan was present, we observed a loop closing motion resembling that shown in Fig. S12. While interactions with LLP and L-tryptophan are certainly expected to be relevant upon the large loop reaching its fully closed state and establishing canonical contacts with these molecules, our results shown above demonstrate that the initial loop closing motion is largely decoupled from the chemical details of the coenzyme and the substrate.

#### Supplementary Note 3

The model of the closed-state *Cr*TDC was constructed by superimposing the open *Cr*TDC structure onto the closed conformation of *Ps*TyDC and subsequently threading the *Cr*TDC loops on the *Ps*TyDC structure. The differences between the MD results and the open *Cr*TDC model were measured via RMSD calculations.

**Supplementary Table 1.**
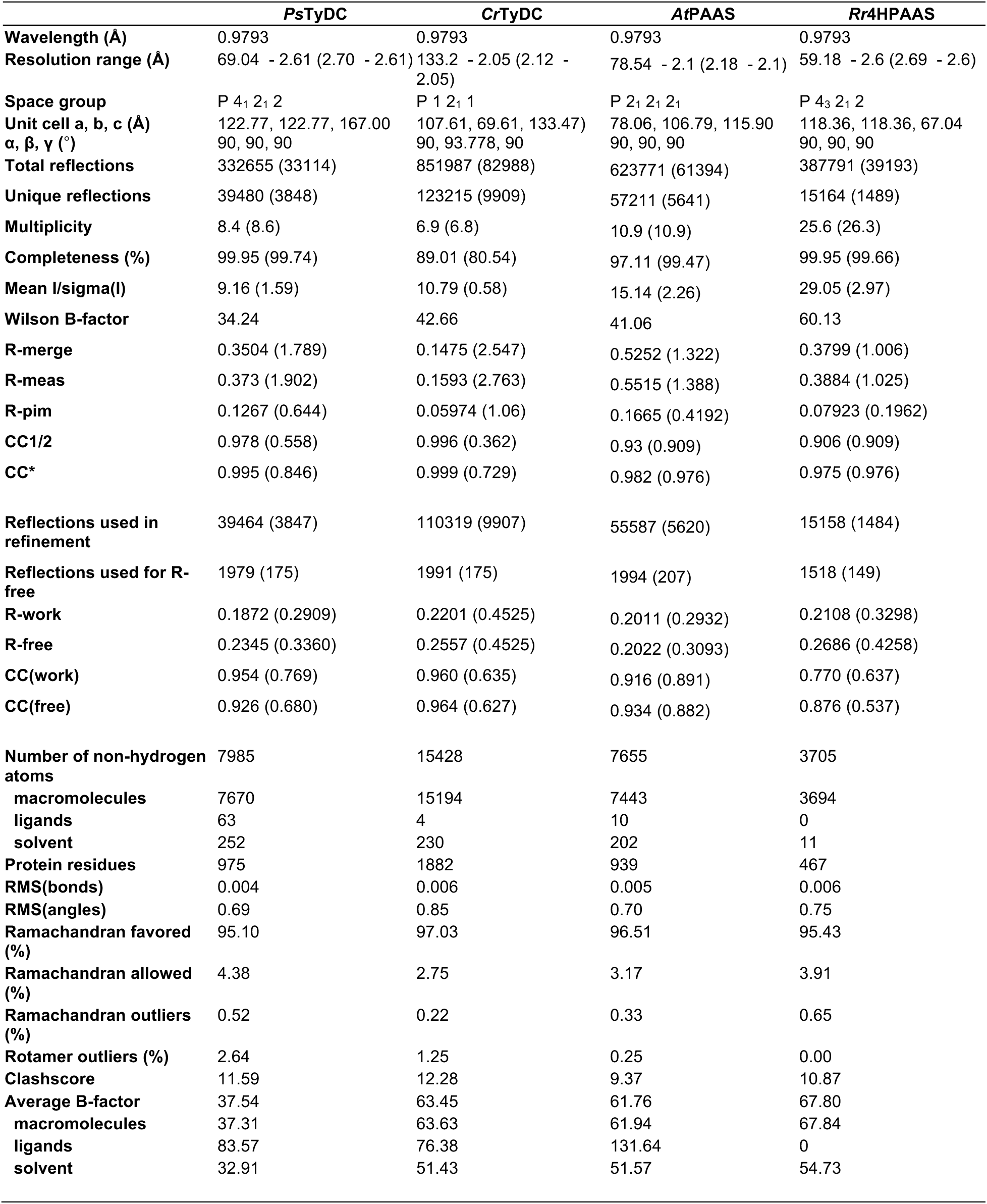
Data collection and structure refinement statistics. Statistics for the highest-resolution shell are shown in parentheses.

|  | *PsTyDC* | *CrTyDC* | *AtPAAS* | *Rr4HPAAS* |
| --- | --- | --- | --- | --- |
| **Wavelength (Å)** | 0.9793 | 0.9793 | 0.9793 | 0.9793 |
| **Resolution range (Å)** | 69.04 - 2.61 (2.70 - 2.61) | 133.2 - 2.05 (2.12 - 2.05) | 78.54 - 2.1 (2.18 - 2.1) | 59.18 - 2.6 (2.69 - 2.6) |
| **Space group** | P 4<sub>1</sub> 2<sub>1</sub> 2 | P 1 2<sub>1</sub> 1 | P 2<sub>1</sub> 2<sub>1</sub> 2<sub>1</sub> | P 4<sub>3</sub> 2<sub>1</sub> 2 |
| **Unit cell a, b, c (Å)** | 122.77, 122.77, 167.00 | 107.61, 69.61, 133.47) | 78.06, 106.79, 115.90 | 118.36, 118.36, 67.04 |
| **α, β, γ (°)** | 90, 90, 90 | 90, 93.778, 90 | 90, 90, 90 | 90, 90, 90 |
| **Total reflections** | 332655 (33114) | 851987 (82988) | 623771 (61394) | 387791 (39193) |
| **Unique reflections** | 39480 (3848) | 123215 (9909) | 57211 (5641) | 15164 (1489) |
| **Multiplicity** | 8.4 (8.6) | 6.9 (6.8) | 10.9 (10.9) | 25.6 (26.3) |
| **Completeness (%)** | 99.95 (99.74) | 89.01 (80.54) | 97.11 (99.47) | 99.95 (99.66) |
| **Mean I/sigma(I)** | 9.16 (1.59) | 10.79 (0.58) | 15.14 (2.26) | 29.05 (2.97) |
| **Wilson B-factor** | 34.24 | 42.66 | 41.06 | 60.13 |
| **R-merge** | 0.3504 (1.789) | 0.1475 (2.547) | 0.5252 (1.322) | 0.3799 (1.006) |
| **R-meas** | 0.373 (1.902) | 0.1593 (2.763) | 0.5515 (1.388) | 0.3884 (1.025) |
| **R-pim** | 0.1267 (0.644) | 0.05974 (1.06) | 0.1665 (0.4192) | 0.07923 (0.1962) |
| **CC1/2** | 0.978 (0.558) | 0.996 (0.362) | 0.93 (0.909) | 0.906 (0.909) |
| **CC\*** | 0.995 (0.846) | 0.999 (0.729) | 0.982 (0.976) | 0.975 (0.976) |
| **Reflections used in refinement** | 39464 (3847) | 110319 (9907) | 55587 (5620) | 15158 (1484) |
| **Reflections used for R-free** | 1979 (175) | 1991 (175) | 1994 (207) | 1518 (149) |
| **R-work** | 0.1872 (0.2909) | 0.2201 (0.4525) | 0.2011 (0.2932) | 0.2108 (0.3298) |
| **R-free** | 0.2345 (0.3360) | 0.2557 (0.4525) | 0.2022 (0.3093) | 0.2686 (0.4258) |
| **CC(work)** | 0.954 (0.769) | 0.960 (0.635) | 0.916 (0.891) | 0.770 (0.637) |
| **CC(free)** | 0.926 (0.680) | 0.964 (0.627) | 0.934 (0.882) | 0.876 (0.537) |
| **Number of non-hydrogen atoms** | 7985 | 15428 | 7655 | 3705 |
| **macromolecules** | 7670 | 15194 | 7443 | 3694 |
| **ligands** | 63 | 4 | 10 | 0 |
| **solvent** | 252 | 230 | 202 | 11 |
| **Protein residues** | 975 | 1882 | 939 | 467 |
| **RMS(bonds)** | 0.004 | 0.006 | 0.005 | 0.006 |
| **RMS(angles)** | 0.69 | 0.85 | 0.70 | 0.75 |
| **Ramachandran favored (%)** | 95.10 | 97.03 | 96.51 | 95.43 |
| **Ramachandran allowed (%)** | 4.38 | 2.75 | 3.17 | 3.91 |
| **Ramachandran outliers (%)** | 0.52 | 0.22 | 0.33 | 0.65 |
| **Rotamer outliers (%)** | 2.64 | 1.25 | 0.25 | 0.00 |
| **Clashscore** | 11.59 | 12.28 | 9.37 | 10.87 |
| **Average B-factor** | 37.54 | 63.45 | 61.76 | 67.80 |
| **macromolecules** | 37.31 | 63.63 | 61.94 | 67.84 |
| **ligands** | 83.57 | 76.38 | 131.64 | 0 |
| **solvent** | 32.91 | 51.43 | 51.57 | 54.73 |

**Supplementary Table 2.** Pairwise sequence identity between select AAAD proteins and the root-mean-square deviation (RMSD) between their monomeric structures.

**Percent amino acid identity matrix**
|  | *Rr4PHAAS* | *CrTDC* | *AtPAAS* | *PsTyDC* |
| --- | --- | --- | --- | --- |
| *Rr4PHAAS* | 100 | 48.06 | 48.76 | 50.82 |
| *CrTDC* |  | 100 | 52.15 | 57.14 |
| *AtPAAS* |  |  | 100 | 56.94 |
| *PsTyDC* |  |  |  | 100 |

**Monomeric structural RMSD (Å) matrix**
|  | *Rr4PHAAS* | *CrTDC* | *AtPAAS* | *PsTyDC* |
| --- | --- | --- | --- | --- |
| *Rr4PHAAS* | 0 | 0.567119 | 0.808045 | 0.624838 |
| *CrTDC* |  | 0 | 0.509434 | 0.419963 |
| *AtPAAS* |  |  | 0 | 0.430512 |
| *PsTyDC* |  |  |  | 0 |

As measured by Phenix Superpose PDB files

**Supplementary Table 3.**
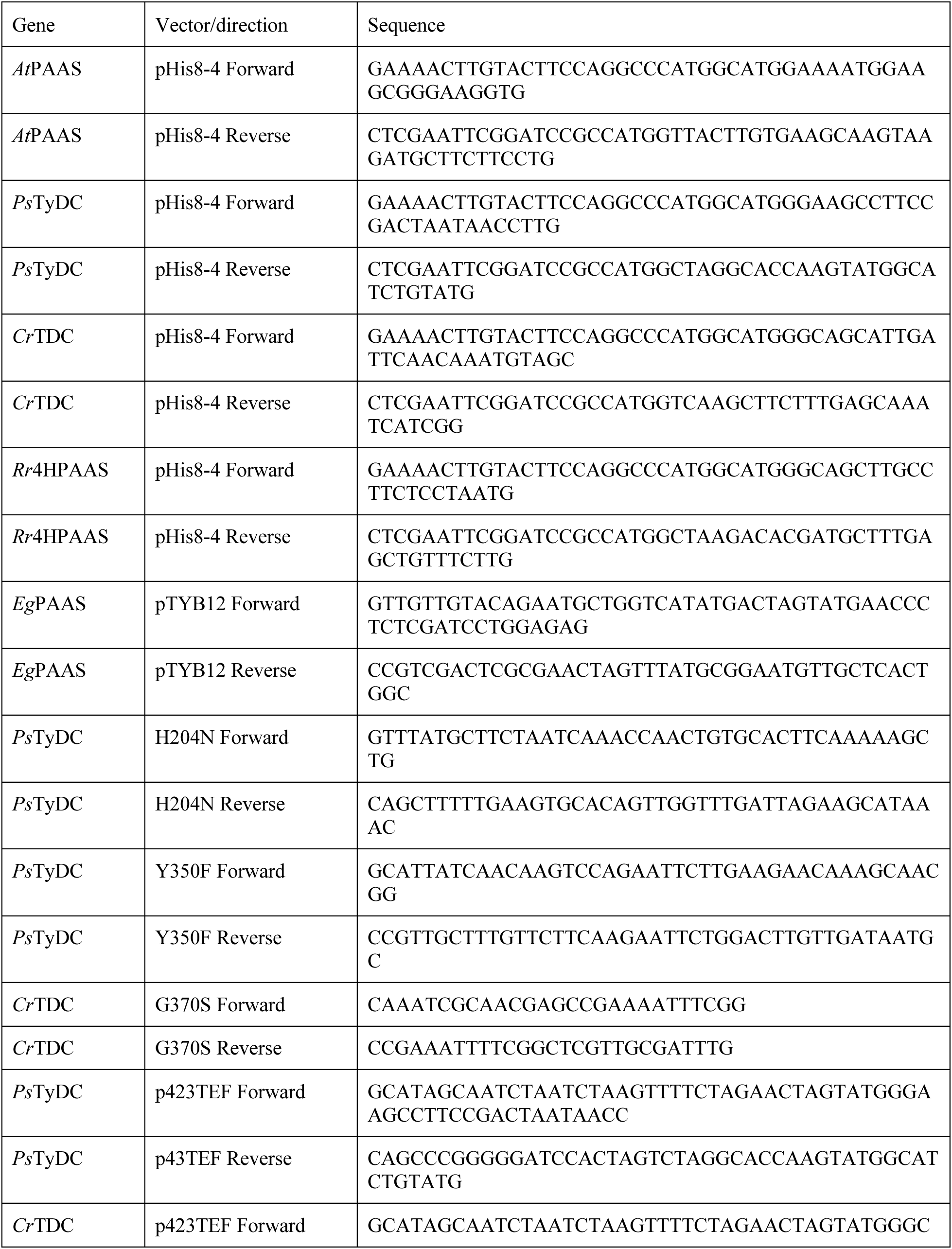
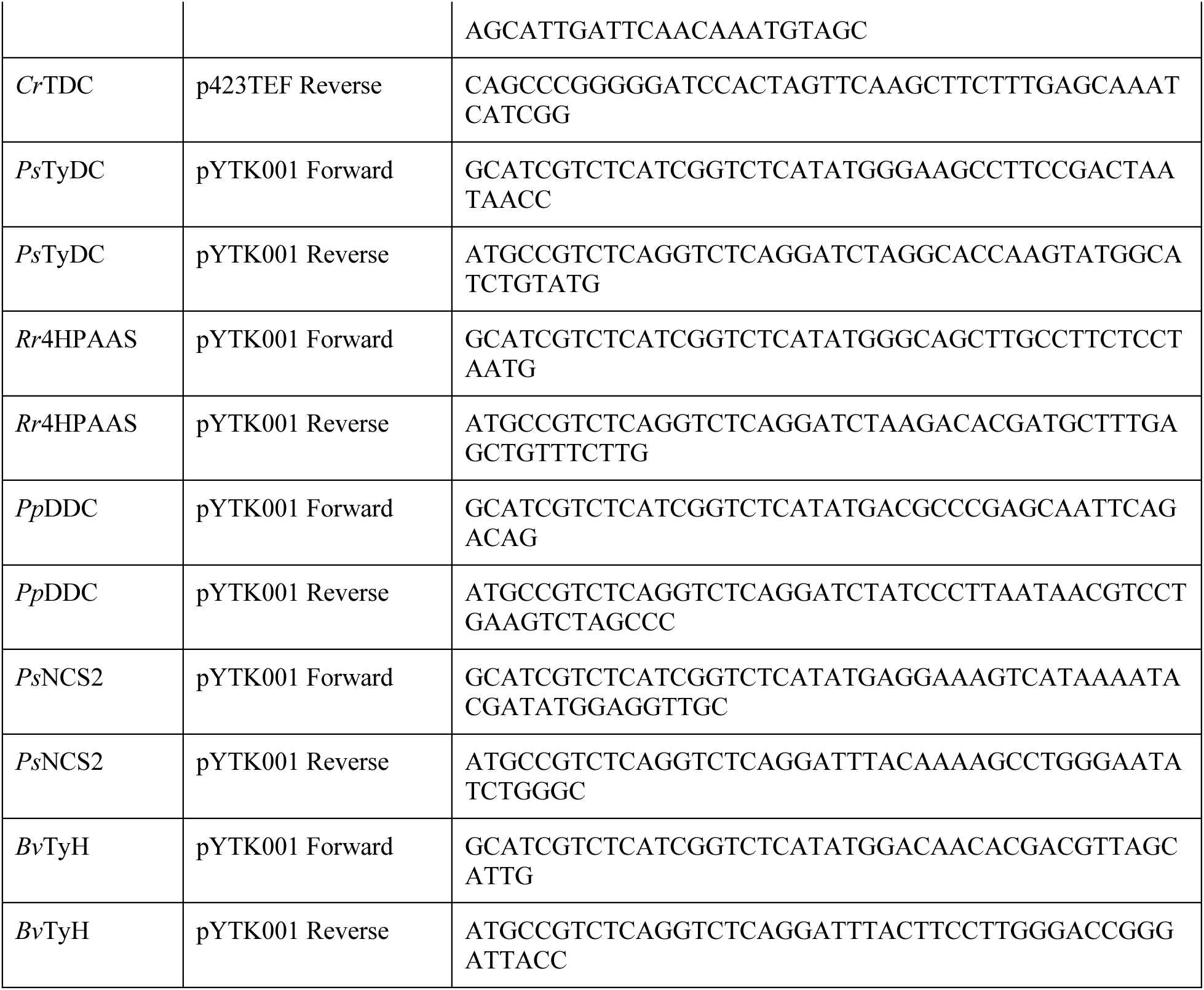
Cloning primers.

**Supplementary Table 4.** List of MD production runs performed in this work. Simulations marked with * were initiated from the end structure of a metadynamics run where the short helix (residues 346-350) on the large loop was forced to unfold (see Methods).

| *Cr*TDC state | LLP and L-tryptophan protonation states | MD simulations performed |
| --- | --- | --- |
| Holo | System 1 | 6 replicas of 100-ns runs with one run extended to 550 ns |
|  |  | 12 replicas of 50-ns runs* |
|  | System 2 | 6 replicas of 100-ns runs |
|  |  | 12 replicas of 50-ns runs* |
|  | System 3 | 6 replicas of 100-ns runs |
|  |  | 12 replicas of 50-ns runs* |
|  | System 4 | 6 replicas of 100-ns runs |
|  |  | 12 replicas of 50-ns runs* |
|  | System 5 | 6 replicas of 100-ns runs |
|  |  | 12 replicas of 50-ns runs* |
|  | System 6 | 6 replicas of 100-ns runs |
|  |  | 12 replicas of 50-ns runs* |
| Apo | - | 3 replicas of 600-ns runs |
| Aggregated simulation time | | ~9.5 μs |

